# Broad-Spectrum Activity and Mechanisms of Action of SQ109 on a Variety of Fungi

**DOI:** 10.1101/2025.02.03.636131

**Authors:** Satish R. Malwal, Rocio Garcia-Rubio, Milena Kordalewska, Hoja Patterson, Chi Zhang, Jorge D. Calderin, Ruijie Zhou, Akanksha M. Pandey, Erika Shor, David S. Perlin, Nathan P. Wiederhold, Luis Ostrosky-Zeichner, Rutilio Fratti, Carol Nacy, Eric Oldfield

**Affiliations:** Department of Chemistry, University of Illinois at Urbana-Champaign, Urbana, Illinois 61801, United States; Center for Discovery and Innovation, Hackensack Meridian Health, Nutley, NJ 07110, United States; Department of Medical Sciences, Hackensack Meridian School of Medicine, Nutley, NJ 07110, United States; Fungus Testing Laboratory, Department of Pathology and Laboratory Medicine, University of Texas Health Science Center at San Antonio, San Antonio, TX 78229, United States; Department of Biochemistry, University of Illinois at Urbana-Champaign, Urbana, Illinois 61801, United States; Department of Microbiology and Immunology, Georgetown University, Washington, DC 20007, United States; Division of Infectious Diseases, University of Texas Health Science Center, Houston, TX 77030, United States; Sequella, Inc., 9610 Medical Center Drive, Suite 200, Rockville, MD 20850, United States

**Keywords:** Antifungals, *Candida*, *Histoplasma*, Protonophore, Vacuoles, Calcium

## Abstract

We investigated the activity of the tuberculosis drug SQ109 against sixteen fungal pathogens: *Candida albicans*, *C. auris*, *C. glabrata*, *C. guilliermondi*, *C. kefyr*, *C. krusei*, *C. lusitaniae*, *Candida parapsilosis*, *C. tropicalis, Cryptococcus neoformans*, *Rhizopus* spp., *Mucor* spp., *Fusarium* spp., *Coccidioides* spp., *Histoplasma capsulatum* and *Aspergillus fumigatus*. MIC values varied widely (125 ng/mL to >64 µg/mL) but in many cases we found promising (MIC∼4 µg/mL) activity as well as MFC/MIC ratios of ∼2. SQ109 metabolites were inactive. The activity of 12 analogs of SQ109 against *Saccharomyces cerevisiae* correlated with protonophore uncoupling activity, suggesting mitochondrial targeting, consistent with the observation that growth inhibition was rescued by agents which inhibit ROS species accumulation. SQ109 disrupted H^+^/Ca^2+^ homeostasis in *S. cerevisiae* vacuoles, and there was synergy (FICI∼0.31) with pitavastatin, indicating involvement of isoprenoid biosynthesis pathway inhibition. SQ109 is, therefore, a potential antifungal agent with multi-target activity.

---

Invasive fungal infections represent a critical and growing health threat that carries significant morbidity and mortality, as well as a substantial economic burden.^1^ The alarming rise in antifungal drug resistance and the emergence of new highly drug-resistant fungal species infecting humans highlight an urgent need for new antifungal drugs.^2^ One approach towards this goal is to repurpose existing drugs used to treat other diseases.^3^ SQ109, a 1,2-ethylenediamine (**1**, Figure 1) developed by Sequella, is a new antitubercular drug candidate. At present, the mechanism of action of SQ109 against yeasts and filamentous fungi has not been reported. However, it is known to target the proton motive force in bacteria (e.g. *Mycobacterium tuberculosis*)^4, 5^ as well as in protozoa, e.g. *Trypanosoma cruzi*,^6^ where it collapses the mitochondrial membrane potential, in addition to disrupting Ca^2+^ homeostasis and affecting sterol biosynthesis.^7^ It also has activity against the bacterium *Helicobacter pylori*.^7^ In mycobacteria, proposed SQ109 targets have included MmpL3 (Mycobacterial membrane protein Large 3; the trehalose monomycolate transporter),^8^ MenA and MenG, involved in menaquinone biosynthesis, as well as decaprenyl diphosphate phosphatase (DPPP)^9, 10^ involved in peptidoglycan biosynthesis. MmpL3, MenA, MenG and DPPP are absent in yeasts/fungi, so are not SQ109 targets, but isoprenoid biosynthesis might be a target, as would uncoupling activity, and based on work with other eukaryotes, calcium homeostasis.^11^

**Figure 1.**
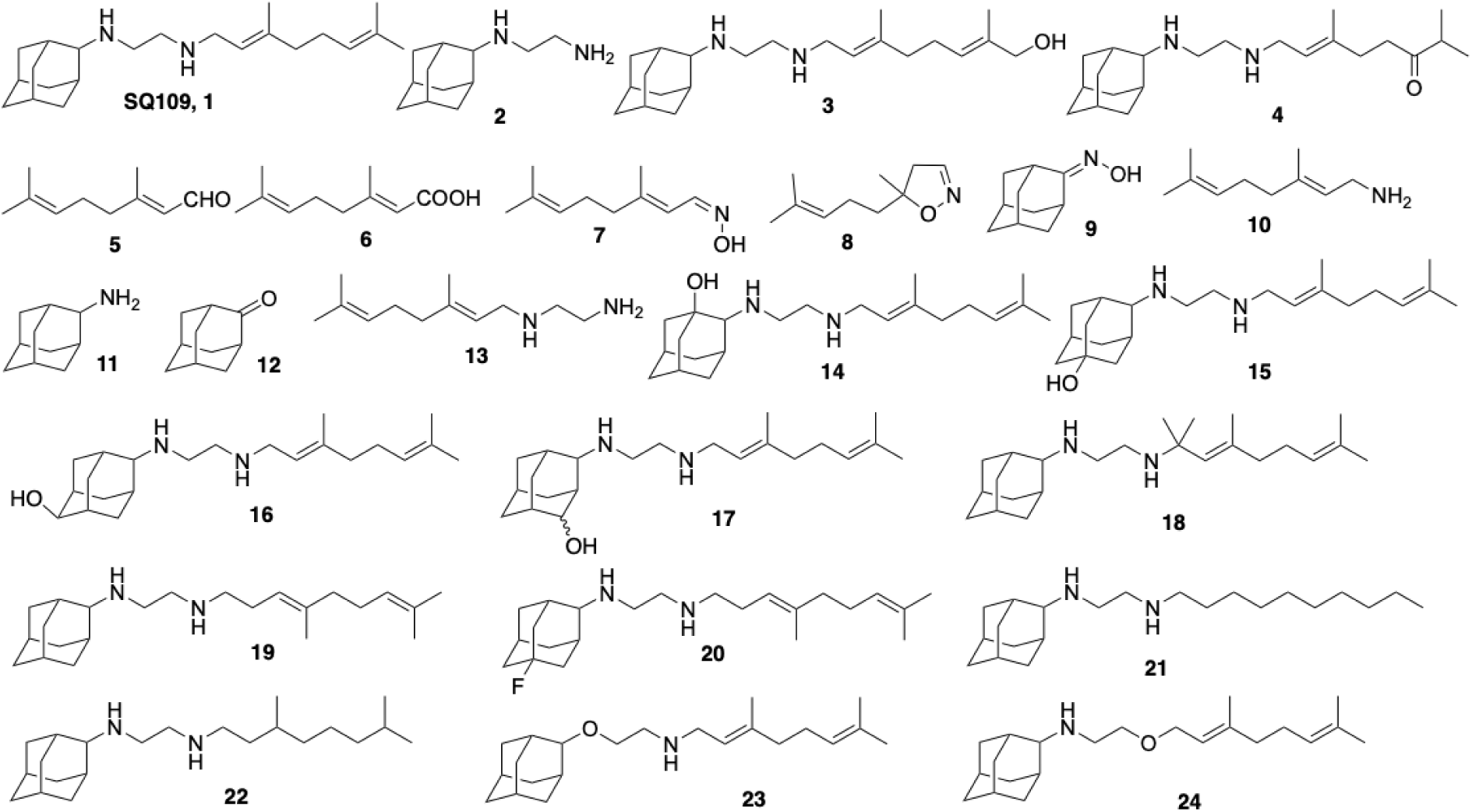
Structures of SQ109, its metabolites and analogs discussed in the Text.

In previous work, Sacksteder et al.^12^ reported that SQ109 had an MIC of 8 μg/mL against *Aspergillus fumigatus* and 1-8 μg/mL against *Candida* spp., and in other work, Li et al.^4^ reported that SQ109 had a 3.3 μg/mL IC_50_ against *Candida albicans* and a 1.1 μg/mL IC_50_ against *Saccharomyces cerevisiae.* Here, we tested SQ109 against 16 different pathogenic fungal species: *C. parapsilosis, C. glabrata, C. krusei, C. tropicalis, C. auris, C. albicans, C. guilliermondi, C. kefyr, C. lusitaniae, Cryptococcus neoformans, Rhizopus* spp*., Mucor* spp*., Fusarium* spp*., Coccidioides* spp*., Histoplasma capsulatum* and *A. fumigatus.* In addition, since SQ109 is known to be metabolized in liver cells^13, 14^ as well as in *M. tuberculosis*^15^, we investigated the 15 known or proposed metabolites^14^ shown in Figure 1, for possible activity since, in principle, SQ109 might be metabolized to a more active species inside fungal cells. We also investigated the possible mechanism of action of SQ109 in the yeast, *S. cerevisiae*, in addition to testing 12 analogs of SQ109 for activity. To probe the mechanistic basis for SQ109 anti-fungal activity, we investigated its effect on H^+^/Ca^2+^ homeostasis in *S. cerevisiae* vacuoles, analogs to the acidocalcisomes found in trypanosomatid and apicomplexan parasites, as well as the activity as protonophore uncouplers for SQ109 and the series of analogs. We also investigated whether reactive oxygen species (from mitochondria) were involved in cell killing, as well as whether the isoprenoid pathway might play a role in activity, using a statin that inhibits isoprenoid biosynthesis.

## RESULTS AND DISCUSSION

### Early investigations of SQ109 activity against yeast and filamentous fungi

During its early preclinical development as a tuberculosis drug candidate, the antifungal activity of SQ109 was evaluated at Sequella and in two independent laboratories with drug-susceptible and drug-resistant *C. albicans,* laboratory and clinical strains of *C. parapsilosis, C. krusei, C. guilliermondii, C. kefyr, C. lusitaniae, C. neoformans, C. tropicalis,* as well as *Aspergillus fumigatus.* At Sequella, we investigated the susceptibility (MIC at 48 hours) of laboratory strains of the yeasts *C. albicans*, *C. glabrata* and *Cryptococcus neoformans* together with the filamentous fungus *Aspergillus fumigatus,* using broth microdilution and following NCCLS (National Committee for Clinical Laboratory Standards, now Clinical and Laboratory Standards Institute) recommended standard methods (M27 and M38 for yeasts and filamentous fungi, respectively).^10, 16^ SQ109 demonstrated activity (8-16 μg/mL) against *C. albicans* and *A. fumigatus,* and more promising activity (0.25-4 μg/mL) against C*. glabrata,* and *Cryptococcus neoformans*, Table S1.

A second set of experiments (Tables S2 and S3) was then carried out under contract to Eurofins/USA (formerly, Focus Bio-Inova) against twenty-four clinical isolates of *C. albicans* (Table S2) that included at least eleven isolates resistant to fluconazole, and *C. parapsilosis* and *C. krusei* (Table S3). The study was again carried out using the broth microdilution method in accordance with CLSI-defined methodology (M27) with amphotericin B as a control, Tables S2 and S3. SQ109 had modest activity against all clinical isolates of *C. albicans*, as well as against *C. parapsilosis*, but better activity against *C. krusei* (1-2 μg/mL). In a third set of experiments, carried out at the University of Texas Health Science Center, Houston, SQ109 was tested against clinical strains of *C. albicans, C. glabrata, C. parapsilosis, C. krusei, C. guilliermondii, C. kefyr, C. lusitaniae, C. tropicalis* and *Cryptococcus neoformans*. Results are shown in Table 1. These results showed that SQ109 had even greater activity against the *C. albicans* strains tested previously and had an MIC of 1-2 µg/mL against several clinical isolates. In addition, SQ109 was found to be effective against *C. parapsilosis, C. krusei, C. guilliermondii, C. kefyr, C. lusitaniae, C. tropicalis* and *Cryptococcus neoformans* with in most cases, MIC values in the ∼1-4 μg/mL range, Table 1.

**Table 1.** Antifungal susceptibility testing of SQ109 against *Candida* spp. and *Cryptococcus neoformans*.

| Organism | # of strains | SQ109MIC (µg/mL) | Organism | # of strains | SQ109MIC (µg/mL) |
| --- | --- | --- | --- | --- | --- |
| *C. albicans* | 3 | 8 | *C. lusitaniae* | 1 | 16 |
|  | 3 | 4 |  | 1 | 2 |
|  | 3 | 2 |  | 3 | 1 |
|  | 1 | 1 |  | 3 | 0.5 |
| *C. glabrata* | 3 | 4 |  | 1 | 0.125 |
|  | 4 | 2 | *Cryptococcus neoformans* | 3 | 2 |
|  | 2 | 1 |  | 3 | 1 |
| *C. guilliermondii* | 1 | 4 |  | 1 | 0.5 |
|  | 4 | 2 | *C. parapsilosis* | 4 | 8 |
|  | 3 | 1 |  | 4 | 4 |
|  | 2 | 0.5 |  | 1 | 2 |
| *C. kefyr* | 1 | 0.25 |  | 1 | 1 |
|  | 4 | 0.125 | *C. tropicalis* | 4 | 4 |
| *C. krusei* | 1 | 2 |  | 3 | 2 |

We then determined the MICs as well as the minimum fungicidal concentrations (MFCs) for SQ109, basically as described previously for *C. albicans*^17^ and *Trichosporon* spp.,^18^ using two strains each of nine yeast species. Results are shown in Table 2 and indicate that SQ109 has a MFC in the 2-8 μg/mL range in 11/18 cases and the average MFC/MIC for all 9 species (18 strains total) is 2.5, indicating fungicidal activity for almost all species.

**Table 2.**
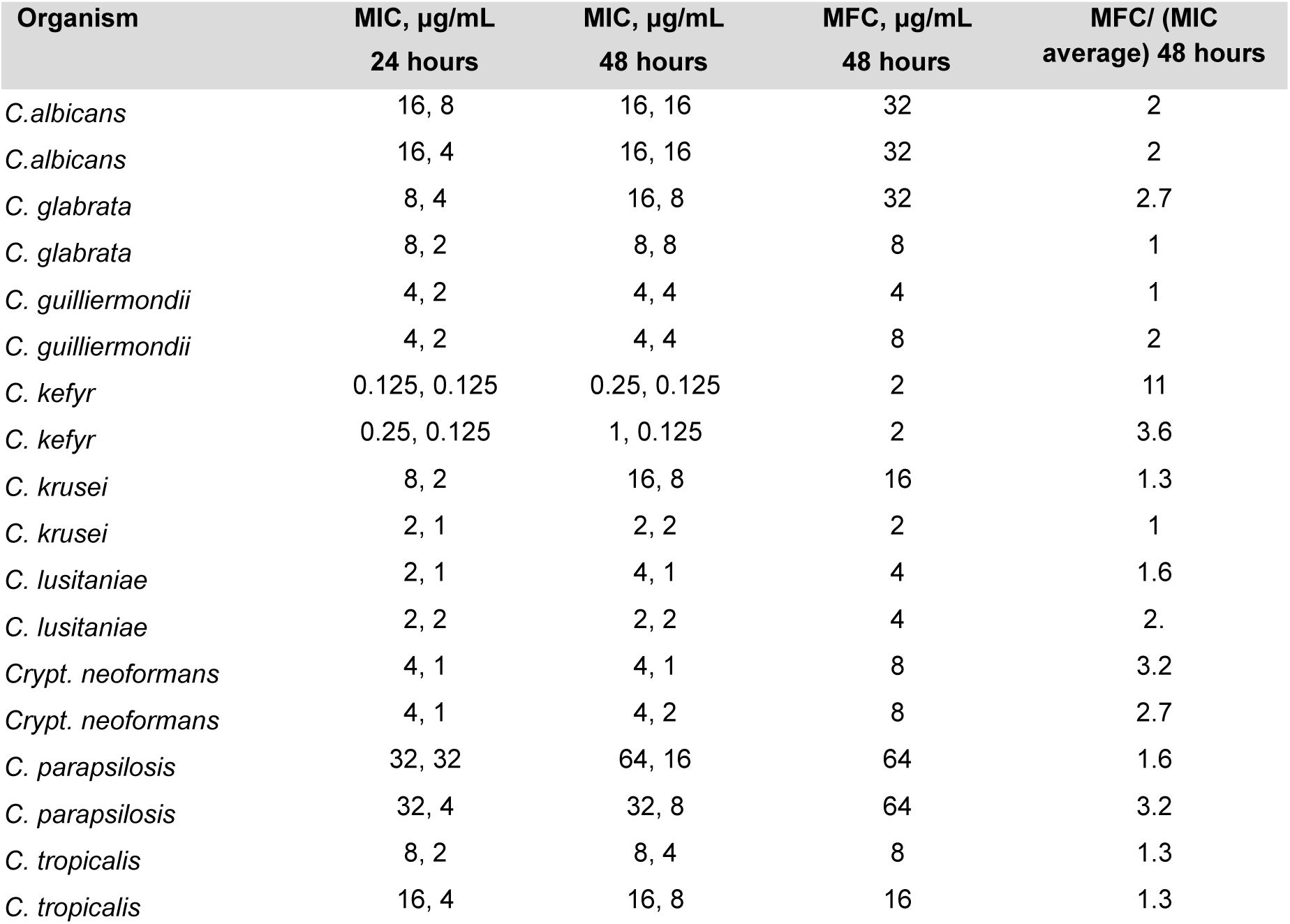
MFC and MIC of SQ109 against *Candida* spp. and *Cryptococcus neoformans*.

In summary, these early results demonstrate activity of SQ109 against clinical strains of *C. albicans* (including fluconazole-resistant strains, MIC 1-8 µg/mL), *C. glabrata* (MIC 0.25-4 µg/mL), *C. parapsilosis* (MIC 1-8 µg/mL)*, C. krusei* (MIC 1-2 µg/mL), *C. guilliermondii* (MIC 0.5-4 µg/mL), *C. kefyr* (MIC 0.125-0.25 µg/mL), *C. lusitaniae* (MIC 0.125-16 µg/mL), *Cryptococcus neoformans* (MIC 0.5-2 µg/mL)*, C. tropicalis* (MIC 1-4 µg/mL), and *A. fumigatus* (MIC 8-16 µg/mL).

### SQ109 activity against azole-susceptible and -resistant yeasts, filamentous fungi, and dimorphic fungi

Given the above results, we expanded our investigation to additional species and strains of fungi, taking into account their azole susceptibility status. Results obtained for *C. albicans, C. auris, C. glabrata,* and *Cryptococcus neoformans* are shown in Table 3 and are of interest since for 25 out of 26 strains, the SQ109 MIC was lower than the fluconazole MIC. Additional results for SQ109 and amphotericin B against 24 clinical strains of *C. albicans* read at 24- and 48-hours of incubation are shown in Table S4 and indicate that the MIC for SQ109 is 8 μg/mL for 21/24 strains at 24 hours and 16 μg/mL at 48 hours, while amphotericin B had an MIC of 1 μg/mL in almost all strains at both 24- and 48-hours.

**Table 3.**
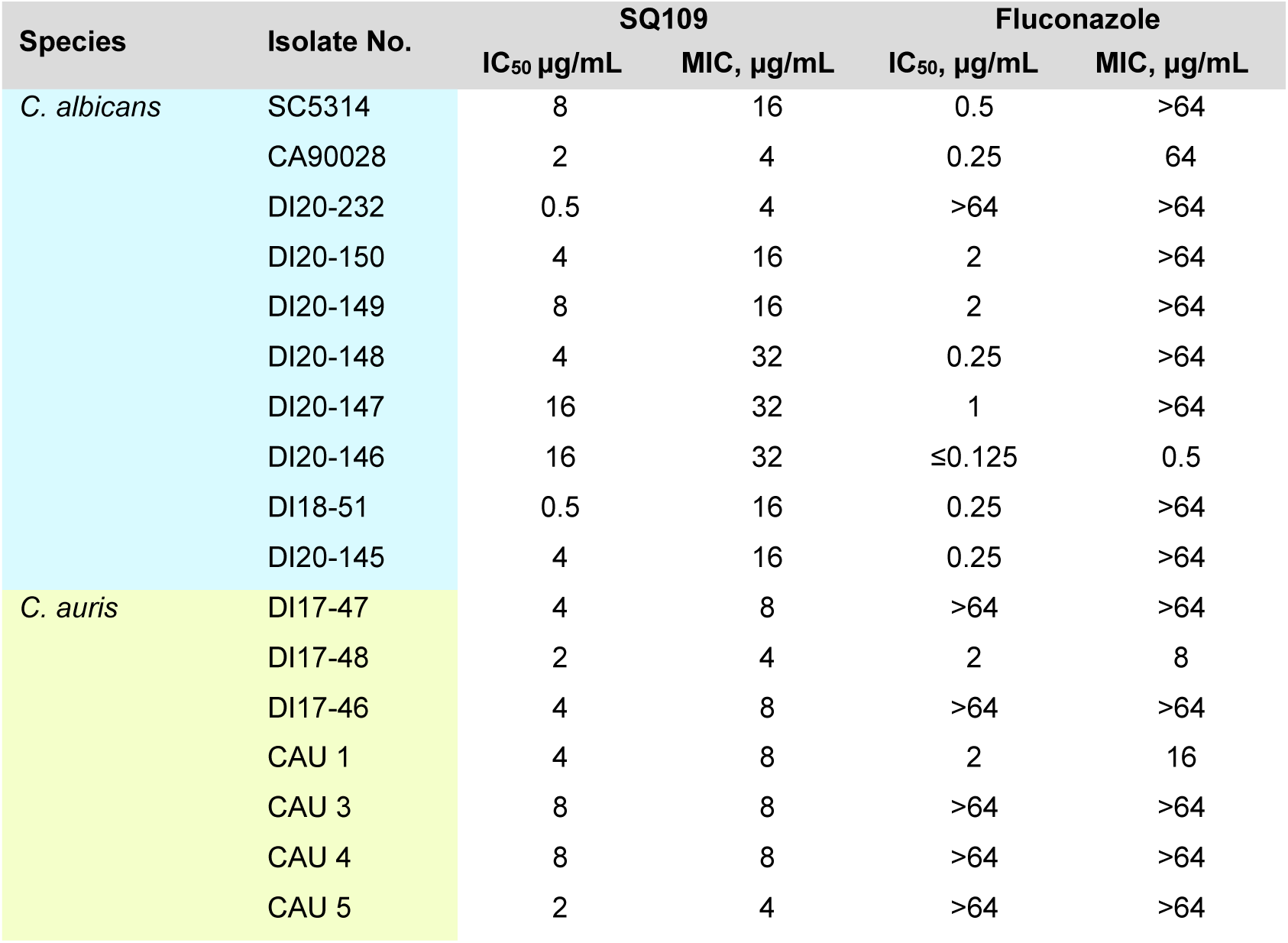
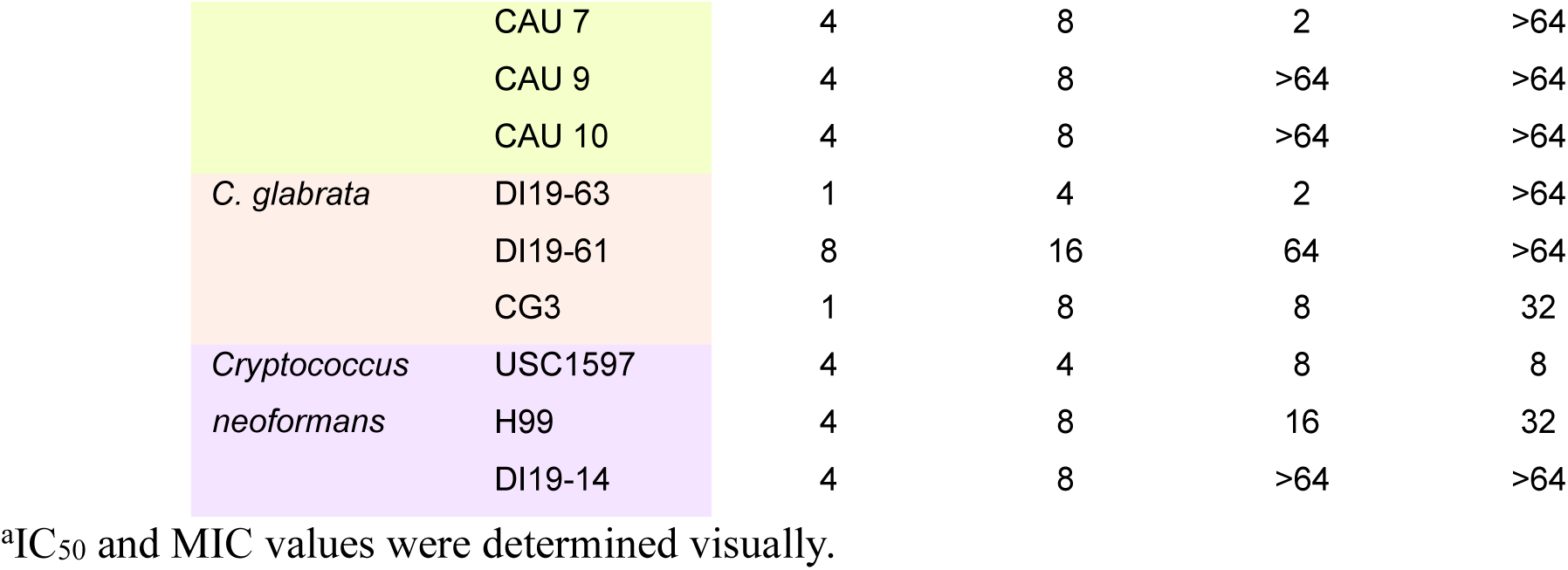
Activity of SQ109 and fluconazole against clinical isolates of *C. albicans, C. auris, C. glabrata* and *Cryptococcus neoformans.*^a^.

SQ109 MICs for clinical isolates of *Rhizopus* spp., *Mucor* spp., *A. fumigatus*, and *Fusarium* spp., filamentous fungi, all showed low activity, ranging from 8-32 μg/mL, as shown in Table 4, consistent with the earlier results we found for *A. fumigatus* (8-16 μg/mL; Table S1).

**Table 4.** Activity of SQ109, posaconazole and/or voriconazole against clinical isolates of the molds *Rhizopus* spp*., Mucor* spp*., Aspergillus fumigatus and Fusarium* spp.^a^.

| Species | Isolate No. | SQ109 |  | Posaconazole |  | Voriconazole |  |
| --- | --- | --- | --- | --- | --- | --- | --- |
|  |  | IC<sub>50</sub> µg/mL | MIC, µg/mL | IC<sub>50</sub>, µg/mL | MIC, µg/mL | IC<sub>50</sub> µg/mL | MIC, µg/mL |
| *Rhizopus* spp. | 99-880 | 16 | 32 | 1 | 2 | --- | --- |
|  | 99-892 | 16 | 32 | 1 | 2 | --- | --- |
|  | DI13-1798 | 8 | 32 | 0.25 | 0.5 | --- | --- |
| *Mucor* spp. | DI15-126 | 8 | 32 | 1 | 4 | --- | --- |
|  | DI15-131 | 8 | 16 | 0.5 | 1 | --- | --- |
|  | DI15-128 | 16 | 32 | 1 | 2 | --- | --- |
| *Aspergillus fumigatus* | AF293 | 16 | 16 | --- | --- | 0.5 | 1 |
|  | DI15-106 | 16 | 32 | --- | --- | >16 | >16 |
|  | DI15-116 | 32 | 32 | --- | --- | 4 | 8 |
|  | AF4 | 32 | 32 | --- | --- | 0.25 | 0.5 |
|  | AF5 | 32 | 32 | --- | --- | 0.5 | 0.5 |
|  | AF6 | 32 | 32 | --- | --- | 0.25 | 0.5 |
|  | AF7 | 32 | 32 | --- | --- | 0.5 | 0.5 |
|  | AF8 | 16 | 32 | --- | --- | 0.5 | 1 |
|  | AF9 | 16 | 16 | --- | --- | 0.5 | 0.5 |
|  | AF10 | 32 | 32 | --- | --- | 2 | 4 |
| *Fusarium* spp. | DI20-73 | 16 | 32 | --- | --- | 4 | 8 |
|  | DI20-76 | 32 | 32 | --- | --- | 0.5 | 8 |
|  | DI19-299 | 8 | 16 | --- | --- | 16 | >16 |

<sup>a</sup>IC<sub>50</sub> and MIC values were determined visually.

However, SQ109 exhibited potent activity against the dimorphic fungi, *Coccidioides* spp. and *Histoplasma capsulatum.* While *Coccidioides* is dimorphic, its two forms are mold and spherule, not the typical mold and yeast forms seen in other dimorphic fungi, such as *Histoplasma*. That notwithstanding, activity against both species was high with MICs ranging from ≤0.125 to 0.25 μg/mL, as shown in Table 5. Activity with fluconazole against *Coccidioides* spp. was poor and voriconazole was not tested, but as expected,^19^ it had good activity against *H. capsulatum*.

**Table 5.**
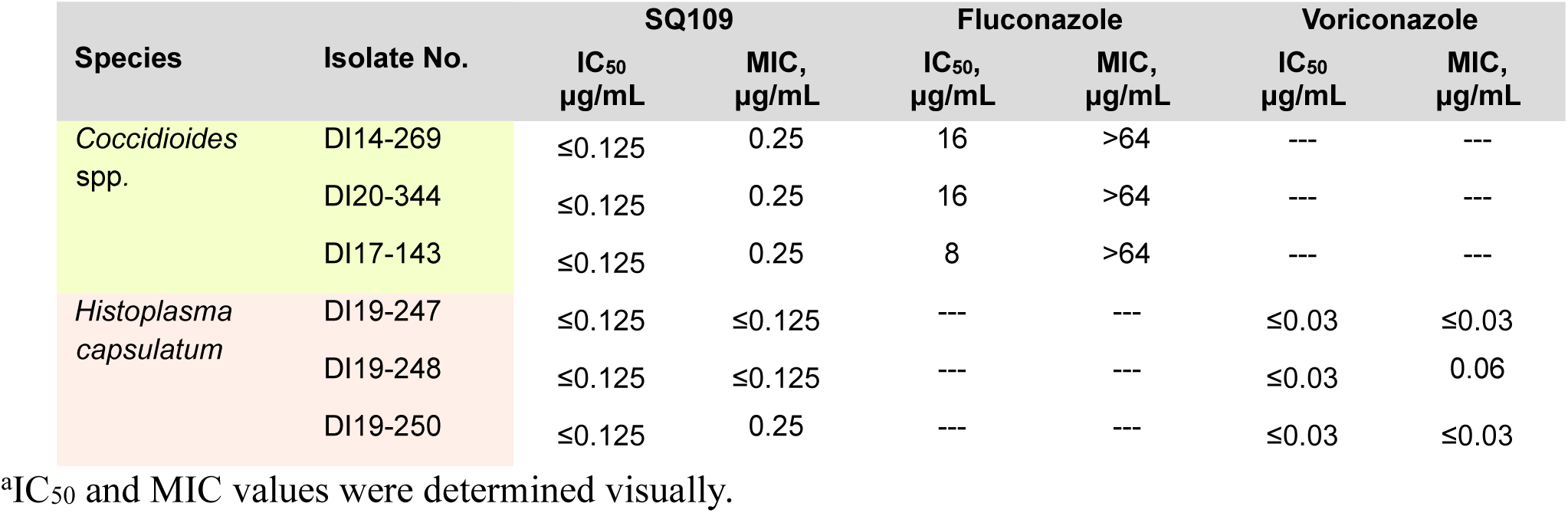
Activity of SQ109, posaconazole or voriconazole against clinical isolates of the dimorphic fungi *Coccidioides* spp*., Histoplasma capsulatum.*^a^.

| Species | Isolate No. | SQ109 |  | Fluconazole |  | Voriconazole |  |
| --- | --- | --- | --- | --- | --- | --- | --- |
| | | IC<sub>50</sub><br>$\mu\text{g/mL}$ | MIC,<br>$\mu\text{g/mL}$ | IC<sub>50</sub>,<br>$\mu\text{g/mL}$ | MIC,<br>$\mu\text{g/mL}$ | IC<sub>50</sub><br>$\mu\text{g/mL}$ | MIC,<br>$\mu\text{g/mL}$ |
| *Coccidioides* spp. | DI14-269 | $\leq 0.125$ | 0.25 | 16 | >64 | --- | --- |
| | DI20-344 | $\leq 0.125$ | 0.25 | 16 | >64 | --- | --- |
| | DI17-143 | $\leq 0.125$ | 0.25 | 8 | >64 | --- | --- |
| *Histoplasma capsulatum* | DI19-247 | $\leq 0.125$ | $\leq 0.125$ | --- | --- | $\leq 0.03$ | $\leq 0.03$ |
| | DI19-248 | $\leq 0.125$ | $\leq 0.125$ | --- | --- | $\leq 0.03$ | 0.06 |
| | DI19-250 | $\leq 0.125$ | 0.25 | --- | --- | $\leq 0.03$ | $\leq 0.03$ |

<sup>a</sup>IC<sub>50</sub> and MIC values were determined visually.

The results with the *Histoplasma* and the *Coccidioides* spp. are of interest for several reasons. First, the IC_50_ values are ≤0.125 µg/mL and the MIC values are in the range ≤0.125 to 0.250 µg/mL, considerably more potent activity than found with most of the other yeasts and filamentous fungi tested. Second, *H. capsulatum* and *Coccidioides* spp. typically reside within macrophages.^20, 21^ and in recent work it has been found that SQ109 polarizes macrophages to the M1 phenotype, as evidenced by transcriptional up-regulation of M1-specific markers such as IL-6, IL-1β, TNF-α, IFN-γ, i-NOS, and down-regulation of the M2-specific markers TGF-β and arginase, resulting in enhanced immune-based killing of *M. tuberculosis* by SQ109.^22^ SQ109 also promoted Th1 and Th17-immune responses that inhibited the bacillary burden in a murine model of TB. Since the macrophage polarization was found even in the absence of the bacterium it seems likely that such effects may also be seen with the dimorphic fungi, resident in macrophages. Third, *M. tuberculosis*, *Histoplasma*, and *Coccidioides* all cause lung infections and can trigger the formation of granulomas, collections of immune cells (primarily macrophages) that try to wall off the infection.^23, 24^ Of interest here is the report that the SQ109 concentrations in lung granulomas (in mice) are ∼50-100 µg/mL.^25^ The combination of low MIC, high concentration in lung granulomas as well as macrophage polarization to the M1 phenotype suggest that SQ19 might have *in vivo* activity against *Histoplasma* and *Coccidioides* spp.

### Activity of SQ109 and potential metabolites against yeasts and fungal pathogens

We next investigated the activity of SQ109 as well as of its metabolites/potential metabolites (shown in Figure 1) against 8 yeast and fungal pathogens. Results for SQ109 are shown in Table 6 and for the metabolites in Tables S5 and S6. In many cases the species investigated contain mutations in one or both of the following proteins: 1,3-beta-D-glucan-UDP glucosyltransferase (FKS) and lanosterol 14-alpha-demethylase (ERG11, called CYP51A in *A. fumigatus*). The mutations in these proteins lead to drug resistance and results for SQ109 are shown in Table 6. The MIC results shown in Table 6 are of interest for several reasons. First, SQ109 has similar or in some cases slightly better activity against the (known) mutant species investigated. Second, there is promising activity against the newly emergent, highly virulent pathogen *C. auris*^26^ where the MIC values against South Asian, South American, East Asian as well as South African clades are in the 2-8 μg/mL range (at both 24 and 48 hours). Third, the results are all consistent with those discussed in our earlier unpublished work.

**Table 6.** Activity of SQ109 against pathogenic yeasts and a filamentous fungus.

|  |  |  |  |  |  |  |  |
| --- | --- | --- | --- | --- | --- | --- | --- |
| *Candida parapsilosis* | FKS | 24 hr | 48 hr | *Candida auris* | FKS/ERG11 | 24 hr | 48 hr |
|  |  | MIC | MIC |  |  | MIC | MIC |
|  |  | (µg/mL) | (µg/mL) |  |  | (µg/mL) | (µg/mL) |
| DPL1017/<br>ATCC22019 | wt | 16 | 16 | DPL1349<br>(S. Asia) | WT/Y132F | 4 | 4 |
| ActDPL1 | wt | 16 | 16 | DPL1371<br>(S. Asia) | S639F/K143R | 4 | 4 |
| DPL8 | wt | 16 | 16 | DPL1373<br>(S. Asia) | S639F/K143R | 4 | 4 |
| DPL300 | wt | 16 | 32 | DPL1384<br>(S. Asia) | WT/WT | 8 | 8 |
| DPL113 | wt | 16 | 16 | DPL1408<br>(S. America) | WT/K177R, N335S,<br>E343D, Y501H | 2 | 2 |
| DPL289 | wt | 16 | 16 | AR-0381<br>(E. Asia) | WT/WT | 4 | 4 |
| DPL1058 | wt | 16 | 32 | AR-0383<br>(S. Africa) | WT/V126A, F126L | 8 | 8 |
| DPL1077 | wt | 16 | 32 |  |  |  |  |
| *Candida glabrata* | FKS |  |  | *Candida albicans* | FKS |  |  |
| CBS138/<br>ATCC2001 | wt | 2 | 4 | DPL1001 | wt | 16 | 16 |
| DPL23 | 2-F641del | 4 | 8 | DPL1002 | wt | 16 | 16 |
| DPL274 | 2-F659S | 0.25 | 0.5 | DPL12 | F641S | 4 | 8 |
| DPL217 | 2-S663F | 2 | 8 | DPL21 | S645P | 0.25 | 2 |
| DPL236 | 2-L664R | 4 | 8 | DPL1014 | R1361R/H | 4 | 8 |
| DPL38 | 1-F625S | 4 | 8 | DPL1039 | F641S/F + S645S/P | 16 | 16 |
| DPL157 | 1-S629P | 2 | 4 | DPL1040 | R1361H | 4 | 8 |
| *Candida krusei* | FKS |  |  | *Aspergillus fumigatus* | CYP51A |  |  |
| DPL1022/<br>ATCC6258 | wt | 1 | 2 | ATCC<br>204304 | wt | 8 | 16 |
| DPL303 | wt | 1 | 2 | ATCC13073 | wt | 16 | 16 |
| DPL310 | wt | 1 | 2 | AF293 | F46Y/M172V/N248T/<br>D255E/E427K | 16 | 16 |
| DPL311 | wt | 1 | 4 | 102 | TR34/L98H | 16 | 16 |
| DPL319 | wt | 1 | 2 | 96 | TR46/Y121F/T289A | 16 | 16 |
| DPL341 | wt | 1 | 1 | F7075 | G54E | 16 | 16 |
| DPL1289 | wt | 1 | 2 | F12865 | G138C | 16 | 16 |
| CLY16038 | wt | 1 | 2 | AF90 | M220V | 16 | 32 |
| *Candida tropicalis* | FKS |  |  | *Candida tropicalis* | FKS |  |  |
| DPL80 | F641S | 4 | 8 | DPL156 | F641S | 4 | 8 |
| DPL228 | S645S/P | 4 | 8 | DPL1302 | wt | 16 | 32 |
| DPL229 | S645S/P | 4 | 8 | DPL55 | wt | 4 | 8 |
| DPL293 | wt | 8 | 8 |  |  |  |  |

Abbreviations used are: FKS=1,3-beta-D-glucan-UDP glucosyltransferase; ERG11=lanosterol 14-alpha demethylase; CYP51A=cytochrome P450 14-alpha sterol demethylase. The mutations in the respective proteins that lead to resistance are indicated.

We then tested the 16 metabolites (or potential metabolites) shown in Figure 1 that we recently synthesized and tested against *Mycobacterium tuberculosis* and *M. smegmatis*.^15^ These compounds are known or likely SQ109 metabolites and some are formed in the liver^14, 15^, so it was of interest to see if any had anti-fungal activity. Essentially all were inactive, having MIC >64 μg/mL, the exception being the 6’-OH SQ109 analog **17** that had activity against *C. krusei*. Full MIC values for all fungi and all compounds tested are given in the Supporting Information (Tables S5 and S6). The lack of activity of any of the SQ109 metabolites against any of the yeasts or fungi investigated strongly suggests that it is SQ109 and not a metabolite that has activity against these organisms, consistent with previous results with *Mycobacterium tuberculosis*.

### Uncoupling activity and effects of SQ109 on *S. cerevisiae* vacuoles H^+^/Ca^2+^ homeostasis

As noted in the introduction, SQ109 is known to have rapid effects on Ca^2+^ signaling in *T. cruzi, T. brucei, L. mexicana,* and *L. donovani*. SQ109 and its analogs also act as protonophore uncouplers in mycobacteria and we previously showed, using an inverted membrane vesicle (IMV) assay,^4^ that cell growth inhibition in *M. smegmatis* correlated with uncoupling activity^9^ (ΔpH and Δψ collapse). We thus next tested SQ109 and the 12 analogs (**3, 4,** and **14-23)** shown in Figure 1 for activity against *Saccharomyces cerevisiae*, as well as their effects as protonophore uncouplers. The synthesis and characterization of all compounds has been reported previously.^4, 14, 9^

As can be seen in Figure 2, we find a correlation (R=0.70, P=0.0077) between *S. cerevisiae* cell growth inhibition (log IC_50_, μM) and ΔpH collapse in the IMV assay (log IC_50_, μM), Figure S1 and Table S7, consistent with uncoupling contributing to cell growth inhibition, as also found in trypanosomatid parasites,^27, 11^ and mycobacteria.^28, 5^

**Figure 2.**
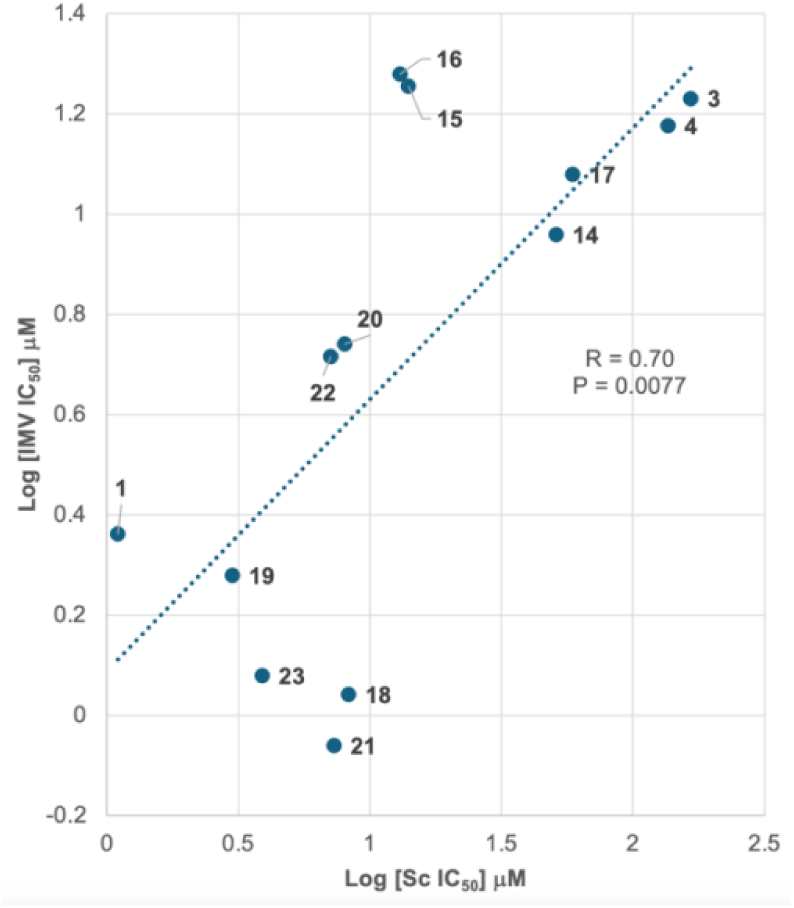
Correlation between *S. cerevisiae* cell growth inhibition by SQ109 and 12 analogs (measured here) and its effect on ΔpH collapse in *E. coli* inverted membrane vesicles, reported previously.

We hypothesized that disruption of mitochondrial function by SQ109 would result in increased formation of reactive oxygen species (ROS). To test that hypothesis, we investigated the inhibition of *S. cerevisiae* cell growth by SQ109 in the presence of five species that inhibit ROS formation: ascorbic acid (AA), *N*-acetyl cysteine (NAC), reduced glutathione (GSH), retinoic acid (RA) and pyrrolidine dithiocarbamate (PDTC). AA, GSH, RA and PDTC have been shown to reverse the growth inhibitory effects of fluconazole against *Cryptococcus neoformans*^29^, due to inhibition of ROS formation. Results for the growth inhibition-rescue of *S. cerevisiae* (ATCC 208352) by AA, RA, GSH, NAC and PDTC are shown in Table 7. The concentrations used are basically those used in the *Cryptococcus neoformans* study.^29^

**Table 7.**
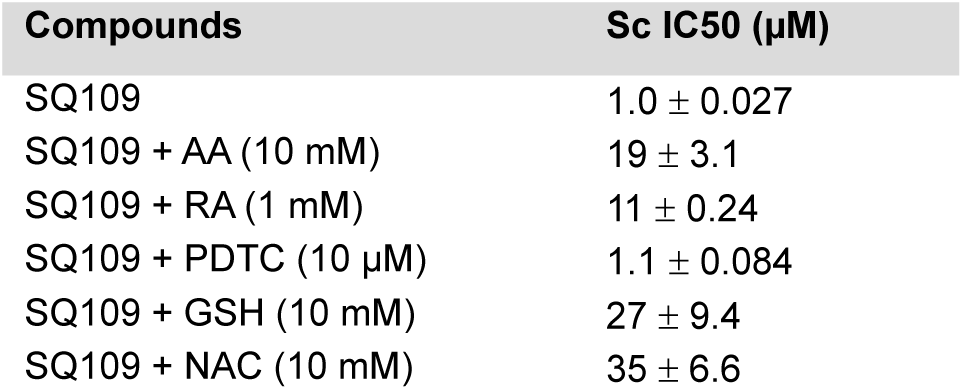
Effects of reactive oxygen species inhibitors on *S. cerevisiae* (ATCC 208352) cell growth inhibition by SQ109.

As can be seen in Table 7, there are on average ∼10x increases in the IC_50_ values on incorporation of AA, RA, GSH and NAC. AA, GSH and NAC are reducing agents, and RA is a polyene likely to scavenge free radicals, so their effects on ROS have a simple chemical basis. PDTC is known for its metal-chelating properties, which can inhibit metal-catalyzed ROS generation, such as the Fenton reaction. However, if the primary source of ROS generated in these yeast cells by SQ109 is not metal-catalyzed, PDTC would be ineffective. Thus, the results shown in Table 7 show that SQ109 is likely to mediate cell killing, again at least in part, via mitochondrial uncoupling leading to increased electron flux and ROS formation.

### Effects of SQ109 on *S. cerevisiae* vacuoles: a calcium connection

SQ109 affects calcium homeostasis in acidocalcisomes in trypanosomatid parasites,^11^ where it increases cytoplasmic calcium concentrations, [Ca^2+^]_in_, and acidocalcisomes have similarities with *S. cerevisiae* vacuoles in their structure and composition.^30^ We thus next investigated the effects of SQ109 on H^+^ and Ca^2+^ transport using isolated *S. cerevisiae* vacuoles. We used a different strain BJ3505 to that discussed above (ATCC 208352), since this was used in our previous work on vacuoles.^31^

We first determined the effects of SQ109 on *S. cerevisiae* BJ3505 growth rate (Figure S2a), as well as the time-dependence of the IC_50_ values for cell growth inhibition (Figure S2b) in order to provide a comparator for experiments on H^+^ and Ca^2+^ transport, in vacuoles.

At 15 hours (where vacuoles were isolated) we found an IC_50_ for growth inhibition of ∼1.5 μM, in good accord with the results on ATCC 208352 discussed above. We then tested SQ109 in the vesicle acidification assay described elsewhere^32^, which measures H^+^ transport catalyzed by the vacuolar (V-type) H^+^-ATPase. Specifically, we used the pH-sensitive fluorophore acridine orange (AO, *N,N,N*′*,N*′-tetramethylacridine-3,6-diamine), which gets trapped in acidic compartments (vacuoles), and its fluorescence decreases as the vacuole is acidified upon adding ATP. We show in Figure 3a a representative real-time experiment tracking AO fluorescence over time. The “ATP” reaction (thick black line) is the positive control, while the DMSO control shows that the solvent used to dissolve SQ109 has no effect on the reporter system. SQ109 was added at either t=0 sec or after a 400 sec incubation. In both cases, we found that SQ109 blocked acidification in a dose-dependent manner. The protonophore uncoupler FCCP (carbonyl cyanide-*p-*trifluoromethoxyphenylhydrazone) was added at the end of the assay to show that AO fluorescence was due to formation of a proton gradient. Average fluorescence (at t=600 sec) for 4 replicates is shown in Figure 3b. As can be seen in Figures 3a and 3b, SQ109 collapses the pH gradient formed by the V-ATPase. We then repeated the experiments, but this time using 8 SQ109 concentrations. Results are shown in Figure 3c and again clearly show that SQ109 collapses the proton gradient and that its effects are similar to those seen with FCCP. Results showing the inhibition of the proton gradient as a function of concentration are shown in Figure 3d and correspond to an IC_50_ of 46 μM. This corresponds to an IC_50_ of 15 μg/mL, which is higher than the IC_50_ for cell growth inhibition. However, for multi-target inhibition, the effects on individual targets will of course be less than seen when all targets are inhibited, plus, drug concentrations in cells and organelles (such as vacuoles and mitochondria) are not known. Next, we tested SQ109 in the Ca^2+^ transport assay reported previously^32^, in which we used the Ca^2+^ sensitive fluorophore Cal520-dextran, which senses Ca^2+^ outside of isolated vacuoles. Ca^2+^ is transported into the vacuole lumen upon adding ATP, leading to a loss in Cal520-dextran fluorescence. Ca^2+^ is taken up until vacuoles start to interact, leading to SNARE (Soluble *N*-ethylmaleimide-sensitive factor Attachment protein Receptors) pair formation between vesicles.^33^ SNARE pairing triggers Ca^2+^ release, which levels off after 20-30 min. This causes Cal520 to become fluorescent. We show in Figure 4a a representative real-time experiment tracking Cal520 fluorescence over time. The “No ATP” control stays flat, showing that no Ca^2+^ is being taken up. The “a**-**Sec17” (antibody against Sec17) trace shows a reaction in which SNARE function is blocked, so Ca^2+^ is taken up, but not released.

**Figure 3.**
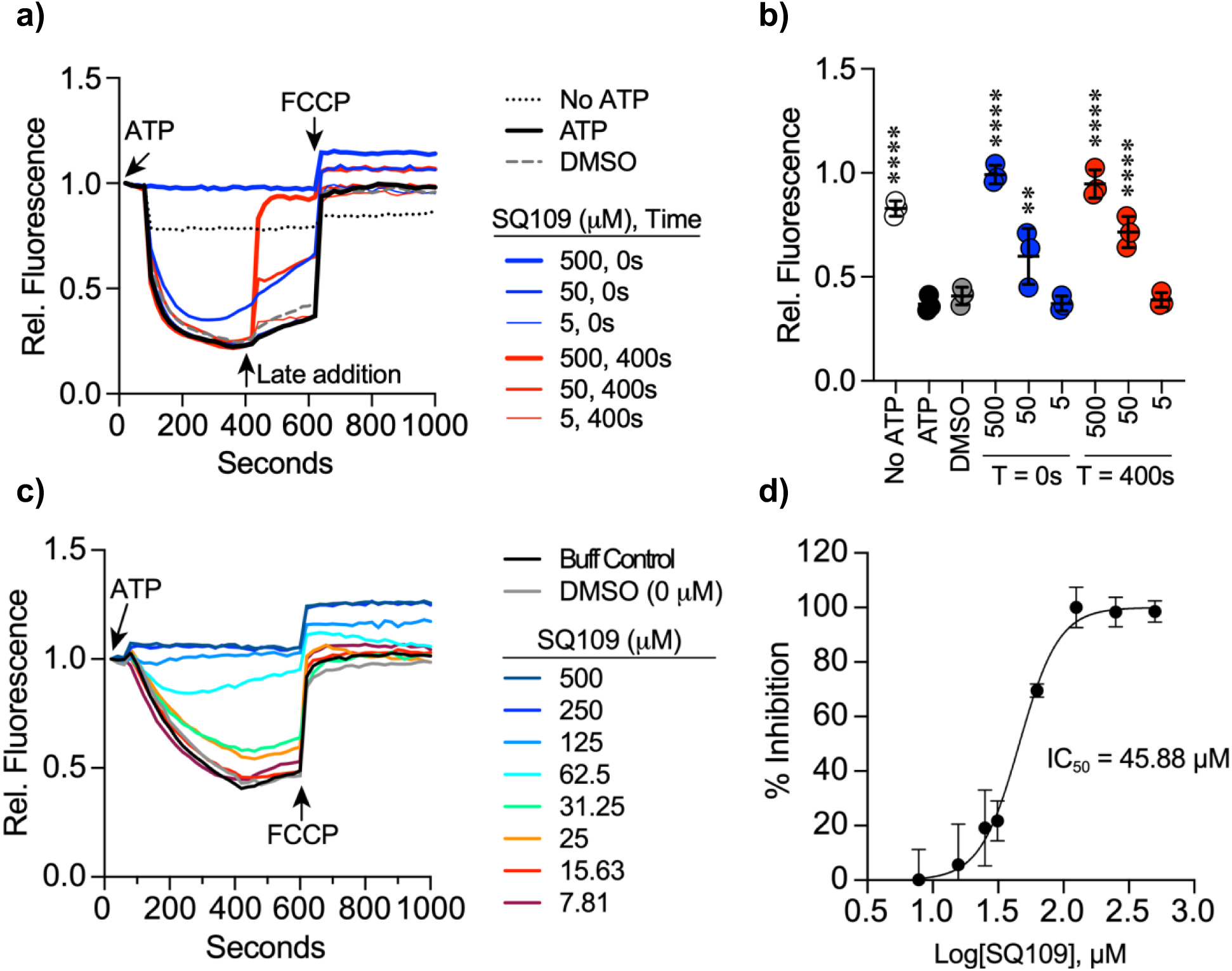
Effects of SQ109 on H^+^ uptake into *S. cerevisiae* (ATCC BJ3505) vacuoles as monitored by acridine orange fluorescence. a) H^+^ uptake with SQ109 added at t=0 and t=400 sec. b) as in a) but data for 4 replicates. Ordinary one-way un-paired ANOVA for multiple comparisons was performed with “ATP” as a control. Error bars are mean +/-SD. Dunnett multiple comparison test was used for individual p values (n=3). *p<0.05, **p<0.01, *** p<0.001, ****p<0.0001. c) H^+^ uptake with SQ109 added at t=0 and for 8 SQ109 concentrations. d) Dose response curve for inhibition of H^+^ uptake by SQ109 for 4 repeat experiments.

**Figure 4.**
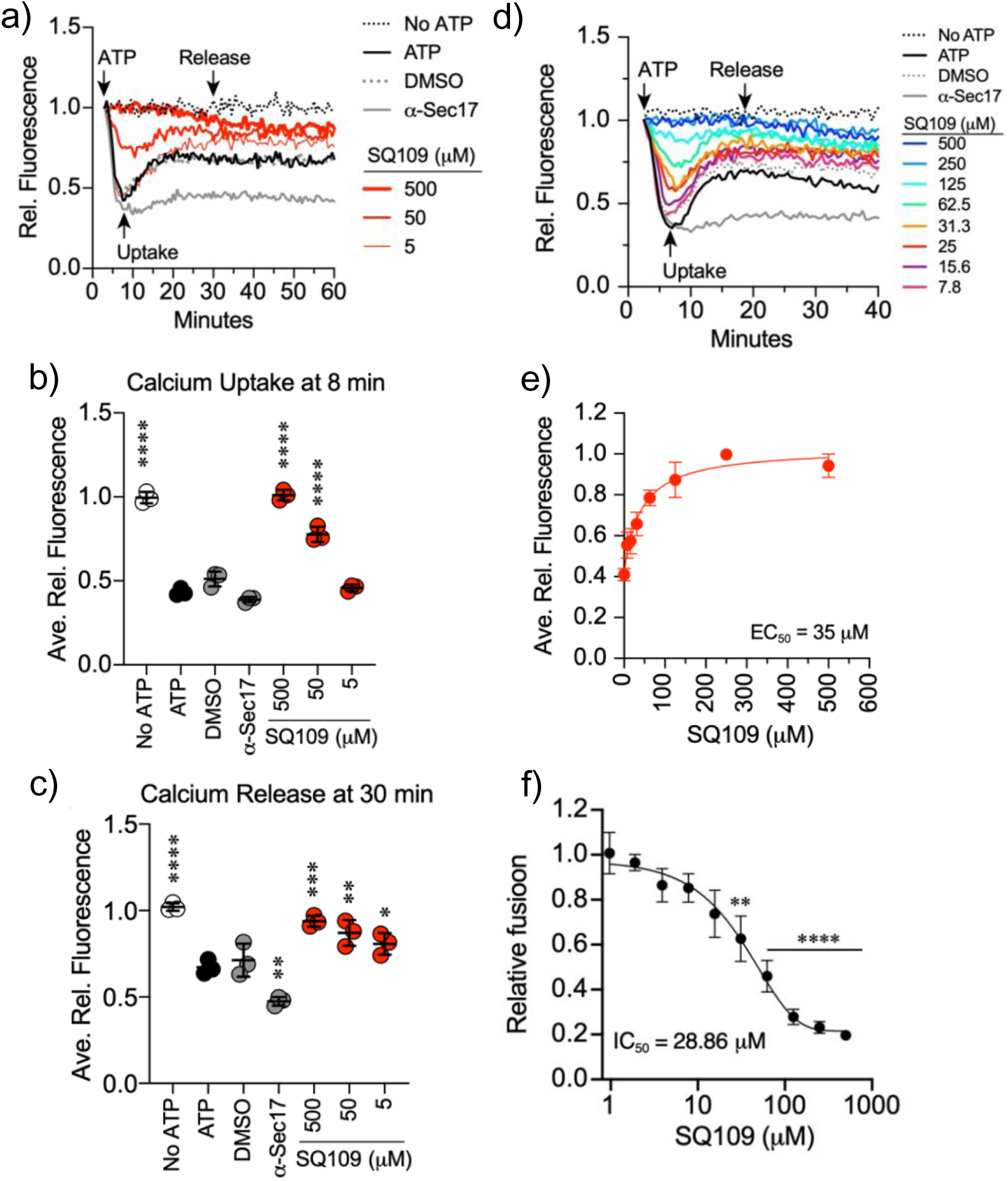
Effects of SQ109 on Ca^2+^ uptake/release in *S. cerevisiae* (ATCC BJ3505) vacuoles as monitored by Cal520-dextran fluorescence, together with vacuole fusion results. a) Ca^2+^ uptake/release at 3 SQ109 concentrations. b) Ca^2+^ uptake at 8 minutes. Ordinary one-way un-paired ANOVA for multiple comparisons was performed with “ATP” as a control. Error bars are mean +/-SD. Dunnett multiple comparison test was used for individual p values (n=3). *p<0.05, **p<0.01, *** p<0.001, ****p<0.0001. c) Ca^2+^ release at 30 minutes. Ordinary one-way un-paired ANOVA for multiple comparisons was performed with “ATP” as a control. Error bars are mean +/-SD. Dunnett multiple comparison test was used for individual p values (n=3). *p<0.05, **p<0.01, *** p<0.001, ****p<0.0001. d) a) Ca^2+^ uptake/release at 8 SQ109 concentrations. SQ109 added at t=0. e) Dose response curve for inhibition of Ca^2+^ uptake. The IC_50_ value is 35 μM. f) Effects of SQ109 on vacuole fusion. The IC_50_ is 29 μM (n=4).

The “ATP” reaction is the positive control and the DMSO reaction shows that the SQ109 solvent has no effect on the reporter system. We found that SQ109 blocked Ca^2+^ uptake in a dose-dependent manner. The average of Ca^2+^ uptake (at 8 min) is shown in Figure 4b, and average peak Ca^2+^ release at 30 min is shown in Figure 4c. We then repeated the Ca^2+^ uptake/release experiment using 8 SQ109 concentrations, Figure 4d, leading to the dose-response curve shown in Figure 4e which has an EC_50_ of 35 μM, similar to that seen with the H^+^-uptake results, Figure 4c. In addition, we measured vacuole fusion directly, finding that SQ109 resulted in deranged vacuole fusion, Figure 4f, with an IC_50_ of 29 μM. Thus, the H^+^-uptake, Ca^2+^ uptake as well as vacuole fusion experiments all indicate significant effects of SQ109 on yeast vacuoles that are expected to contribute to its effect on cell growth inhibition.

### Synergistic effects of SQ109 with a statin

In recent work it was found that the statin drug pitavastatin had activity against *C. albicans* NR-29448 (MIC=8 μM) and that it acted synergistically with the azole drug fluconazole. This strain is resistant to fluconazole (MIC=256 μM) but the combination was highly synergistic, with a FICI=0.05.^34^ Pitavastatin targets the enzyme HMG-CoA (hydroxymethyl-coenzymeA) reductase, at the start of the isoprenoid biosynthesis pathway, while fluconazole targets lanosterol 14α-demethylase, involved in ergosterol biosynthesis. Since SQ109 has been shown to have effects on sterol biosynthesis in trypanosomatid parasites,^6^ we investigated whether there might be synergistic effects between pitavastatin and SQ109, which would improve activity. FICI were determined by following previously published protocols. ^35, 36, 37, 38^

We found an FICI=0.31, Figure 5, indicating strong synergism, consistent with SQ109 acting in the isoprenoid biosynthesis pathway.

**Figure 5.**
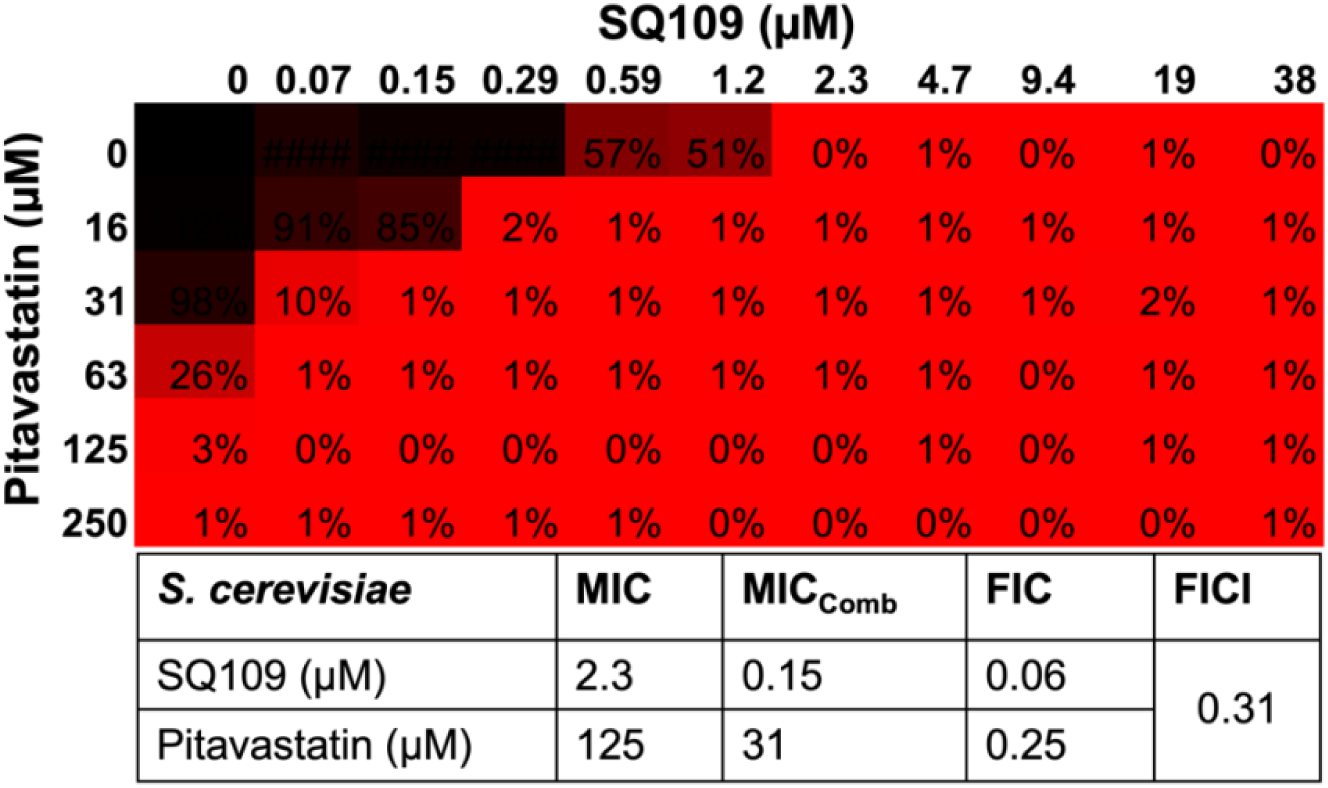
Synergistic effect of SQ109 and pitavastatin in *S. cerevisiae*.

## CONCLUSIONS

The results we have presented above are of interest for several reasons. First, we tested SQ109 for activity against 16 species of pathogenic yeasts and molds, finding promising activity (low μg/mL) values for many of the yeasts, including drug resistant strains. Results were obtained in five independent laboratories, with good agreement between the different groups. SQ109 metabolites were all basically inactive. Second, we began an investigation of the mechanism of action of action of SQ109, against *S. cerevisiae*. We found effects of SQ109 on H^+^ and Ca^2+^ transport in yeast vacuoles, and a correlation between protonophore uncoupling activity and cell growth inhibition for SQ109 and a series of SQ109 analogs. Third, we found that cell growth inhibition activity could be rescued (∼10x increase in IC_50_) by agents that reduce reactive oxygen species activity, as seen with fluconazole, due to reduced oxidative stress, proposed here to be due to mitochondrial targeting. Fourth, we found strong synergistic activity (FICI∼0.31) with the statin drug, pitavastatin, consistent with targeting isoprenoid biosynthesis. Our results are reminiscent of the effects of SQ109 in bacteria (ΔpH, Δψ collapse; isoprenoid biosynthesis) as well as in trypanosomatid parasites (Δψ collapse; Ca^2+^ homeostasis, sterol biosynthesis) and are of general interest since they open up new possibilities for developing SQ109, currently in clinical trials for tuberculosis, as an antifungal agent with broad spectrum activity and multiple-site targeting. *Histoplasma* and *Coccidioides* spp. appear of most interest since the MIC values for SQ109 were the among the lowest observed (≤0.125 to 0.250 µg/mL); they reside in macrophages; macrophages are stimulated to the M1 phenotype by SQ109; infected macrophages reside in lung tissue granulomas, and SQ109 is known to accumulate in lung tissue/granulomas to high (∼50-100 µg/mL) levels.

## EXPERIMENTAL SECTION

### Chemicals

The SQ109 sample was the hydrochloride salt prepared under GLP/GMP conditions and used in clinical testing and was provided by Sequella Inc. and was 99.5 % pure by qNMR. SQ109 analogs and metabolites were from the batches described previously.^15^

### Cell Growth Inhibition Assays

Testing was performed at HMH-CDI using broth microdilution methods as described in CLSI documents M27^39^ and M60^40^. Antifungal susceptibility testing for mold isolates was performed in accordance with the guidelines described in CLSI documents M38^41^ and M61.^42^ In brief, all compounds were prepared as 10 mM stock solutions in DMSO and the range of drug concentrations tested was from 64 µg/mL to 0.125 µg/mL in two-fold serial dilutions. All tested strains were part of the Perlin lab collection (HMH-CDI) and represent a variety of known antifungal resistance mechanisms. Each strain was tested in duplicate and consensus 24- and 48-hour minimal inhibitory concentration (MIC) values for each strain and compound were obtained visually.

Antifungal susceptibility testing at the University of Texas Health Science Center, Houston, was carried out as described previously^17^ following the M27-A method in RPMI 1640 buffered to pH 7.0 with 0.165 M MOPS buffer with readings taken at 24 and 48 h. After 48 h, 20 μL of each clear well was plated on SDA (Sabouraud dextrose agar), and the minimum fungicidal concentration (MFC) is the lowest concentration giving no growth (≥98% killing).

Antifungal susceptibility testing at the University of Texas Health Science Section, San Antonio, was performed by using CLSI broth dilution methods as described in the CLSI M27^39^ and M38^41^ standards. Stock solutions of the investigational agent and comparators were prepared at concentrations 100-times the highest concentration to be tested (e.g., 1600 or 6400 μg/ml) in DMSO. Aliquots of the stock solutions were dispensed into polystyrene tubes and stored at −20°C. The synthetic medium RPMI-1640 (with glutamine, without bicarbonate, and with phenol red) was used. The RPMI was buffered to a pH of 7.0 +/-0.1 at 25°C with 0.165 M MOPS (3-[*N*-morpholino] propanesulfonic acid). For broth microdilution, sterile U-shaped 96-well cell culture plates were used for performing the MIC assays. Dilutions of the DMSO stocks were prepared into RPMI to achieve 2x concentrations (1:50 dilution). After the dilutions of the working 2x investigational antifungal solutions were prepared, 0.1 mL of each concentration was transferred into a pre-specified column of the U-shaped 96-well cell culture plate using sterile pipettes. For *Coccidioides* and *Histoplasma*, polystyrene tubes, not 96 well plates, were used, as recommended in the CLSI M38 standard. 0.01 mL of the 100x stock solutions were transferred to test tubes.

*Candida* isolates were grown on Sabouraud dextrose agar (potato flake agar was used for *Aspergillus*) and cells were collected after an appropriate period of growth for each species being evaluated. The fungi were suspended in sterile distilled water, and the densities of the fungal suspension were read using a spectrophotometer and adjusted to an appropriate optical density specific for each fungal species. The fungal suspension of each isolate was then diluted in RPMI. A sufficient volume of the test inoculum was prepared to directly inoculate 0.1 mL into each test well of the 96-well cell culture plate. Final inoculum ranges are dependent on the fungal species to be tested (e.g., 0.4 x 10^3^ to 5 x 10^3^ cells/mL for *Candida*, *Cryptococcus*, *Coccidioides*, and *Histoplasma*, and 0.4 x 10^4^ to 5 x 10^4^ cells/ml for *Aspergillus*, *Fusarium*, *Mucor*, and *Rhizopus*). Each well of the 96-well cell culture plate containing SQ109 and comparators (0.1 mL) was inoculated on the day of the assay with 0.1 mL of the fungal suspension. Growth control wells contained 0.1 mL of fungal suspension and 0.1 mL of the growth medium without antifungal agents. The media control well contained 0.2 mL of the growth medium. As noted above, for *Coccidioides* and *Histoplasma*, polystyrene tubes, not 96 well plates, were used, as recommended in the CLSI M38 standard, and 0.99 mL of the inoculum was added to each tube to achieve the desired concentration.

The microdilution trays and tubes were incubated at 35°C without agitation. After the appropriate period of incubation (e.g., 24 hours for *Candida*, *Rhizopus*, and *Mucor*; 48 hours for *Aspergillus*, *Coccidioides*, and *Fusarium*; 72 hours for *Cryptococcus*, and 168 hours for *Histoplasma*), MIC and IC_50_ values were determined. The MIC is the minimum concentration that results in no visible cell growth, that is, ∼100% cell growth inhibition, while the IC_50_ is the concentration that produces 50% growth inhibition, as estimated visually. One positive comparator/control was used for yeast and *Coccidioides* (fluconazole) and one for the other species (voriconazole or posaconazole). Appropriate quality control isolates were also included on each day of testing as recommended by CLSI (e.g., *C. parapsilosis* ATCC 22019, *C. krusei* ATCC 6258, and *P. variotii* MYA-3630).

### *S. cerevisiae* cell growth inhibition assay

*S. cerevisiae* (ATCC 208352, grown 48 h at 30 °C) was diluted 50-fold in YPD media and grown to an OD_600_ of ∼0.4, which occurred after about 6.5 hrs. The culture was then diluted 500-fold into fresh YPD medium to give a working solution. This working solution (200 μL) was then transferred into every well in a flat-bottomed 96-well plate except for the first well wells of columns of interest. Inhibitors were added at specific starting concentrations (with a total volume of 300 μL (diluted with working solution) to the first well of the columns. The inhibitors were then sequentially diluted 3-fold across 8 wells; 360 μL of autoclaved Milli-Q water was added to empty well to prevent water evaporation from the plate. Plates were incubated at 30 °C, with shaking at 200 rpm for 48 h. The OD_600_ values were then measured and used to determine cell growth inhibition, by using GraphPad Prism software (version 7.04). Experiments were carried out in duplicate or triplicate. IC_50_ values were determined using a four-parameter variable-slope function in the Prism program.

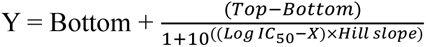

### *S. cerevisiae* cell growth inhibition rescues

These assays were performed following the same procedure as described for *S*. *cerevisiae* cell growth inhibition except that we used twelve 2x serial dilutions. Rescue reagents were added at the concentrations shown in Table 7 (for each well).

### *S. cerevisiae* synergy experiments

Checkerboard assays were performed in 96-well plates following a previously published protocol.^35^ *S. cerevisiae* (ATCC 208352) grown at 30 °C, 200 rpm, for ∼48 h in YPD media, was diluted 50x and grown to an OD_600_ of ∼0.4, followed by a 500x dilution to give a working solution. Pitavastatin and SQ109 in DMSO were added to the plate and were then 2-fold serially-diluted. After incubation at 30 °C, 200 rpm, for ∼48 h, OD_600_ values were then measured, normalized, and a 10% cutoff was assigned to visually determine the MICs. The fractional inhibitory concentration index (FICI) was calculated using the formula^36, 37, 38^: FICI = (MIC_S_^combi^/MIC_S_^alone^) + (MIC_P_^combi^/MIC_P_^alone^), where S=SQ109 amd P=pitavastatin.

### Vacuole proton transport assay

The proton pumping activity of isolated vacuoles was performed as described by others^43, 44^ with some modifications, as described previously.^32^ *In vitro* acidification reactions (60 μL) contained 20 μg vacuoles, reaction buffer (20 mM PIPES-KOH pH 6.8, 200 mM sorbitol, 125 mM KCl, 5 mM MgCl_2_), ATP-regenerating system (1 mM ATP, 0.1 mg/mL creatine kinase, 29 mM creatine phosphate), 10 μM CoA, 283 nM Pbi2 (inhibitor of protease 2), and 15 μM of AO. Reaction mixtures were loaded into a black, half-volume 96-well flat-bottom plate with nonbinding surface. ATP regenerating system or buffer was added, and reactions were incubated at 27 °C while AO fluorescence was monitored. SQ109 was added at the times indicated in Figure 5. Samples were analyzed by using a fluorescence plate reader (POLARstar Omega, BMG Labtech) with the excitation filter at 485 nm and emission filter at 520 nm. Reactions were initiated by the addition of ATP-regenerating system, following the initial measurement. After fluorescence quenching plateaus were reached, 30 μM FCCP was added to collapse the proton gradient and restore AO fluorescence.

### Calcium transport in vacuoles

Vacuolar Ca^2+^ transport was measured as described.^45, 46, 47^ *In vitro* Ca^2+^ transport reactions (60 μL) contained 20 μg vacuoles from BJ3505 backgrounds, reaction buffer, 10 μM CoA, 283 nM Pbi2, and 150 nM of the Ca^2+^ probe Cal-520 dextran conjugate, molecular weight 10,000. Reaction mixtures were loaded into a black, half-volume 96-well flat-bottom plate with nonbinding surface. ATP-regenerating system was added, and reactions were incubated at 27 °C while Cal-520 fluorescence was monitored. Samples lacking ATP were used a negative control for Ca^2+^ uptake. Antibody against the SNARE cochaperone Sec17 (a-Sec17) was added as a negative control for SNARE-dependent Ca^2+^ efflux.^48^ Samples were analyzed using a fluorescence plate reader with the excitation filter at 485 nm and emission filter at 520 nm. Reactions were initiated with the addition of ATP-regenerating system following the initial measurement. The effects of inhibitors on efflux were determined by the addition of buffer or inhibitors immediately following Ca^2+^ influx. Calibration was carried out using buffered Ca^2+^ standards (Invitrogen).

## ASSOCIATED CONTENT

### Supporting Information

The Supporting Information is available free of charge at http://pubs.acs.org.

## Notes

The authors declare no competing financial interests.

## ACKNOWLEDGMENTS

We thank Dr. Sangjin Hong, Department of Biochemistry, University of Illinois at Urbana-Champaign for assistance with the IMV assays.

## ABBREVIATIONS

MmpL3: Mycobacterial membrane protein Large 3
MenA: isoprenyl diphosphate:1,4-dihydroxy-2naphthoate (DHNA) isoprenyltransferase,
MenG: demethylmenaquinone methyl transferase
DPPP: decaprenyl diphosphate phosphatase
ROS: reactive oxygen species
AA: ascorbic acid
RA: retinoic acid
GSH: reduced glutathione
NAC: N-acetyl cysteine
SNARE: SNAP Receptors
SNAP: soluble Nethylmaleimide-sensitive factor attachment proteins (Sec17p in yeast; SQ109 N1-(adamantan-2-yl)-N2[(2E)-3,7-dimethylocta-2,6-dien-1-yl]ethane-1,2-diamine
MIC: minimum inhibitory concentration
MFC: minimum fungicidal concentration
FICI: fractional inhibitory concentration index
FCCP: carbonyl cyanide-p-trifluoromethoxyphenylhydrazone
AO: acridine orange, N,N,N?,N?-tetramethylacridine-3,6-diamine.

