## Supplementary material for "Broad-Spectrum Activity and Mechanisms of Action of SQ109 on a Variety of Fungi": Yeast1 SI_013025.docx

States.

^4^Fungus Testing Laboratory, Department of Pathology and Laboratory Medicine, University of Texas Health Science Center at San Antonio, San Antonio, TX 78229, United States

^5^Department of Biochemistry, University of Illinois at Urbana-Champaign, Urbana, Illinois 61801, United States.

^6^Department of Microbiology and Immunology, Georgetown University, Washington, DC 20007, United States.

^7^Division of Infectious Diseases, University of Texas Health Science Center, Houston, TX 77030, United States.

^8^Sequella, Inc., 9610 Medical Center Drive, Suite 200, Rockville, MD 20850, United States.

**Table of Contents**

| **Figure S1.** Dose response curves for *S. cerevisiae* growth inhibition by SQ109 and analogs. | S3 |
| --- | --- |
| **Figure S2**. Time-dependence of *S. cerevisiae* (ATCC BJ3505) cell growth rate on SQ109 concentration and the time-dependence of the IC_50_. | S4 |
| **Table S1**. Activity of SQ109 against laboratory strains of *Candida albicans*, *Candida glabrata*, and *Aspergillus fumigatus* to SQ109 | S5 |
| **Table S2.** Activity of SQ109 against clinical strains of *C. albicans* | S5 |
| **Table S3.** Activity of SQ109 and amphotericin B against *C. parapsilosis and C. krusei* | S5 |
| **Table S4**. Activity of SQ109 and amphotericin B against 24 clinical strains of *C. albicans* read at 24 and 48 hours of incubation | S5 |
| **Table S5.** Activity of SQ109 and known/potential metabolites against pathogenic fungi | S6 |
| **Table S6.** Activity of SQ109 and known/potential metabolites against pathogenic fungi | S7 |
| **Table S7.** *S. cerevisiae* growth inhibition by SQ109 and analogs, and their *E. coli* inverted membrane vesicles (IMV) IC_50_ | S8 |
| SMILES | S9 |

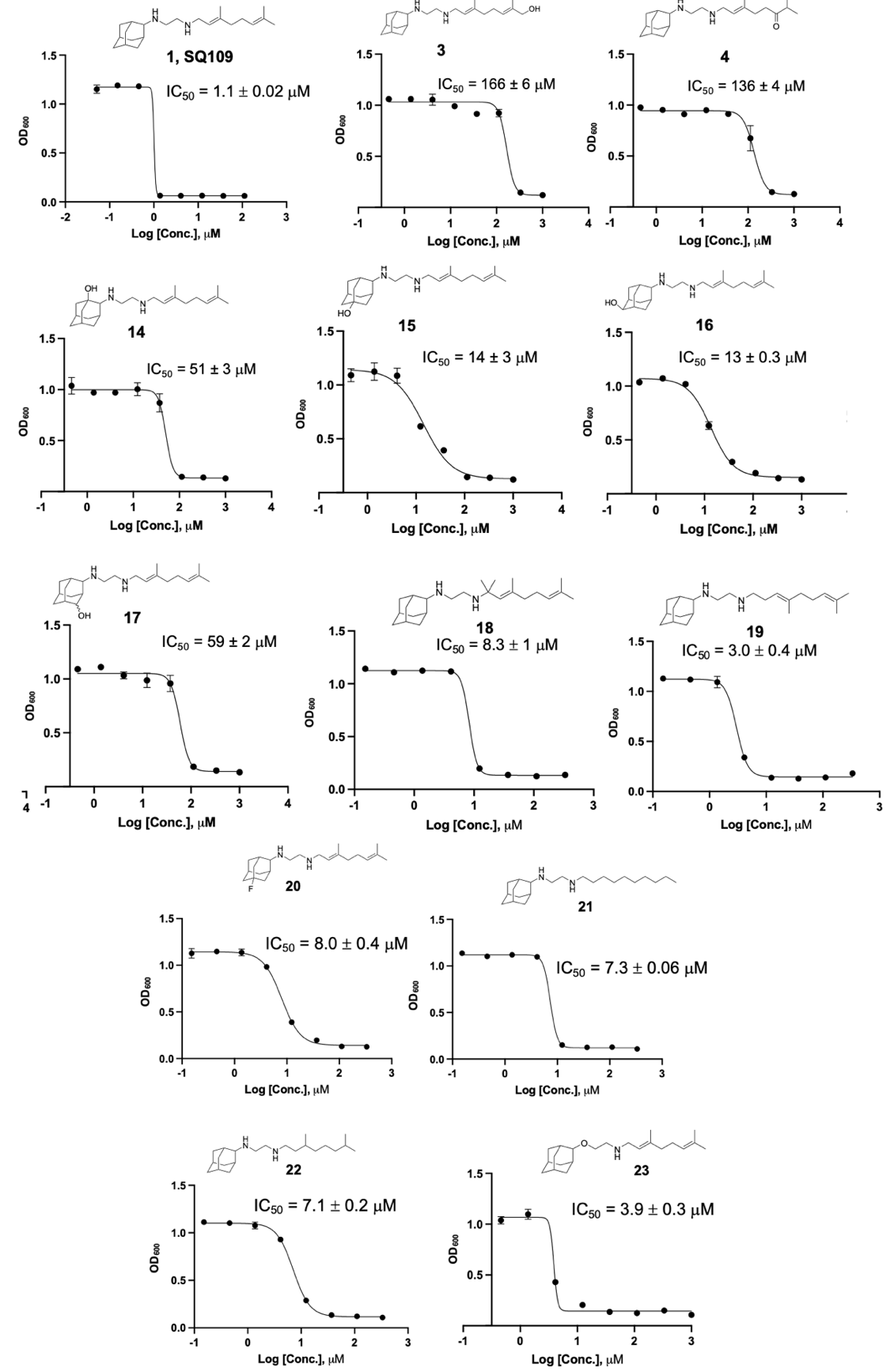

**Figure S1.** *S. cerevisiae* growth inhibition by SQ109 and analogs.

**a)**

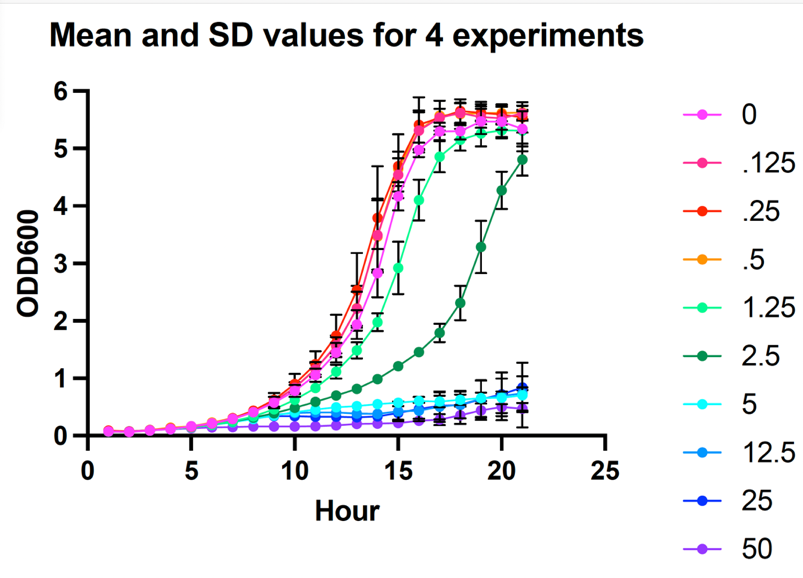

**b)**

| Time (hr) | IC50 mM |
| --- | --- |
| 7 | 2.25 |
| 8 | 2.6 |
| 9 | 2.17 |
| 10 | 1.83 |
| 11 | 1.73 |
| 12 | 1.55 |
| 13 | 1.4 |
| 14 | 1.25 |
| 15 | 1.47 |
| 16 | 1.8 |
| 17 | 2 |
| 18 | 2.18 |
| 19 | 2.78 |
| 20 | 3.2 |
| 21 | 3.38 |

**Figure S2.** a) Time-dependence of *S. cerevisiae* (ATCC BJ3505) cell growth rate on SQ109 concentration. b) Time-dependence of IC_50_ for SQ109 inhibition of *S. cerevisiae* (ATCC BJ3505) cell growth.

**Table S1**. **Activity of SQ109 against laboratory strains of *C. albicans*, *C. glabrata*, and *A. fumigatus***

| **Organism** | **SQ109 MIC (µg/mL)** |
| --- | --- |
| ATCC 90028 *Candida albicans* | 4-8 |
| ATCC 15545 *Candida glabrata* | 0.25-1 |
| ATCC 16424 *Aspergillus fumigatus* | 8-16 |
| *Cryptococcus neoformans* | 0.25-4 |

**Table S2. Activity of SQ109 against clinical strains of *C. albicans***

| **Incubation time** | **Organism** | **Number of Strains** | **MIC (µg/mL)** |
| --- | --- | --- | --- |
| **24 h^1^** | *Candida albicans* | 1 | 16 |
|  | *Candida albicans* | 21 | 8 |
|  | *Candida albicans* | 2 | 4 |
| **48 h^2^** | *Candida albicans* | 2 | 32 |
|  | *Candida albicans* | 17 | 16 |
|  | *Candida albicans* | 4 | 8 |
|  | *Candida albicans* | 1 | 4 |

^1^MIC of amphotericin B at 24 hours, µg/mL: 1 (20 strains), 0.5 (3 strains), 0.25 (1 strain); ^2^ the MIC of amphotericin B at 48 hours was 1 µg/mL for all strains tested.

**Table S3. Activity of SQ109 and amphotericin B against *C. parapsilosis and C. krusei.***

| **Organism** | **SQ109 MIC (µg/mL)** | **Amphotericin B MIC (µg/mL)** |
| --- | --- | --- |
| *Candida parapsilosis* (ATCC 22019) | 8 | 0.5 |
| *Candida parapsilosis* (ATCC 22019) | 16 | 1 |
| *Candida krusei* (ATCC 6258) | 1 | 1 |
| *Candida krusei* (ATCC 6258) | 2 | 2 |

**Table S4**. **Activity of SQ109 and amphotericin B against 24 clinical strains of *C. albicans* read at 24 and 48 hours of incubation; all values in μg/mL**

| **Agent & timepoint** | **Number of MIC values at each concentration** | | | | | | | | | | **Range** | **MIC**  **50/90** | **Mode** |
| --- | --- | --- | --- | --- | --- | --- | --- | --- | --- | --- | --- | --- | --- |
|  | **0.125** | **0.25** | **0.5** | **1** | **2** | **4** | **8** | **16** | **32** | **64** |  |  |  |
| SQ109 24 hrs |  |  |  |  |  | 2 | 21 | 1 |  |  | 4-16 | 8 / 8 | 8 |
| AMB 24 hrs |  | 1 | 3 | 20 |  |  |  |  |  |  | 0.25-1 | 1 / 1 | 1 |
| SQ109 48 hrs |  |  |  |  |  | 1 | 4 | 17 | 2 |  | 4-32 | 16 / 16 | 16 |
| AMB 48 hrs |  |  |  | 24 |  |  |  |  |  |  | 1 | 1 / 1 | 1 |

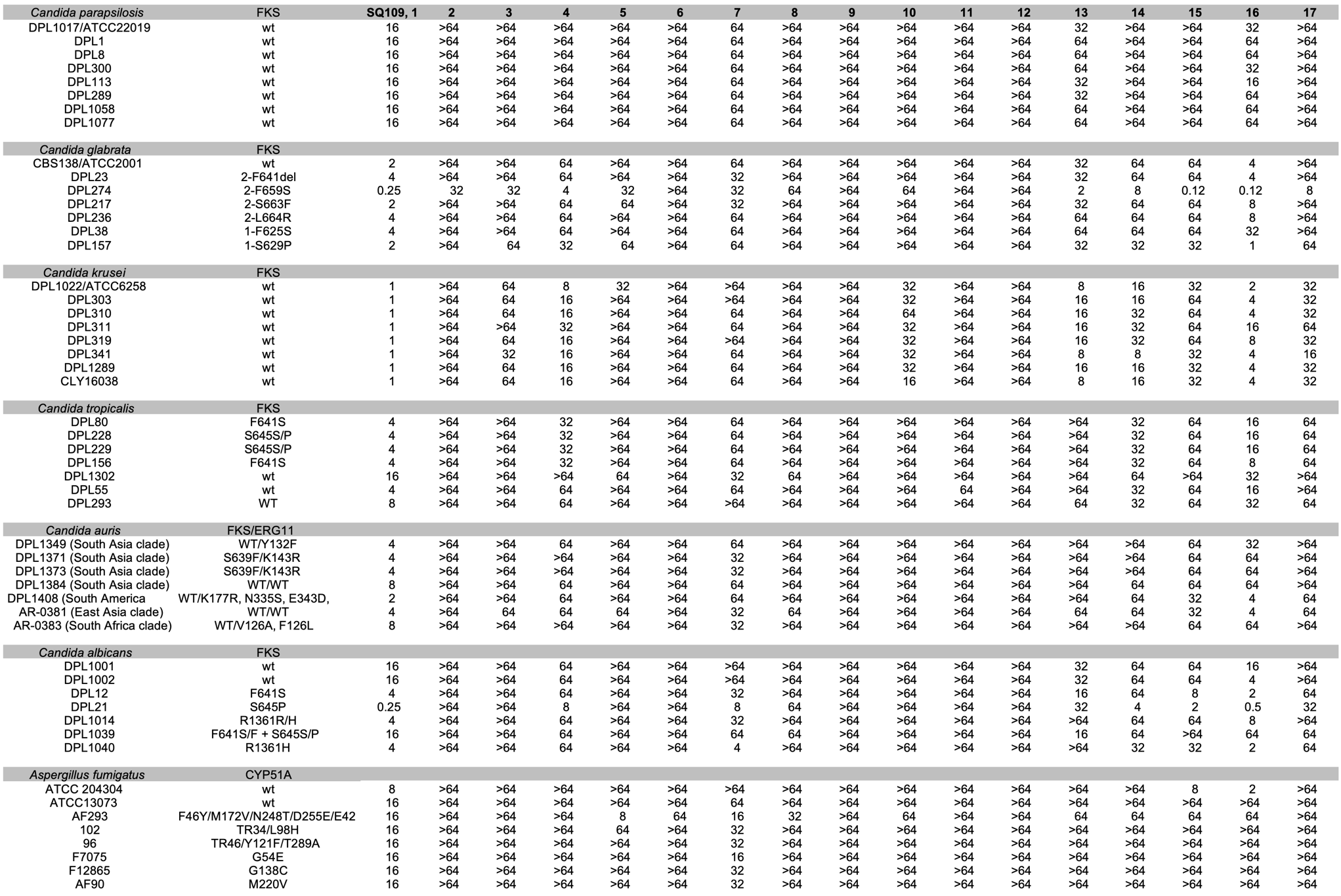

**Table S5. Activity of SQ109 and known/potential metabolites against pathogenic fungi^a^.**

^a^MIC is in mg/mL after 24 h. Abbreviations used are: FKS=1,3-beta-D-glucan-UDP glucosyltransferase; ERG11=lanosterol 14-alpha-demethylase; CYP51A=cytochrome P450 14-alpha sterol demethylase. The mutations in the respective proteins that lead to resistance are indicated.

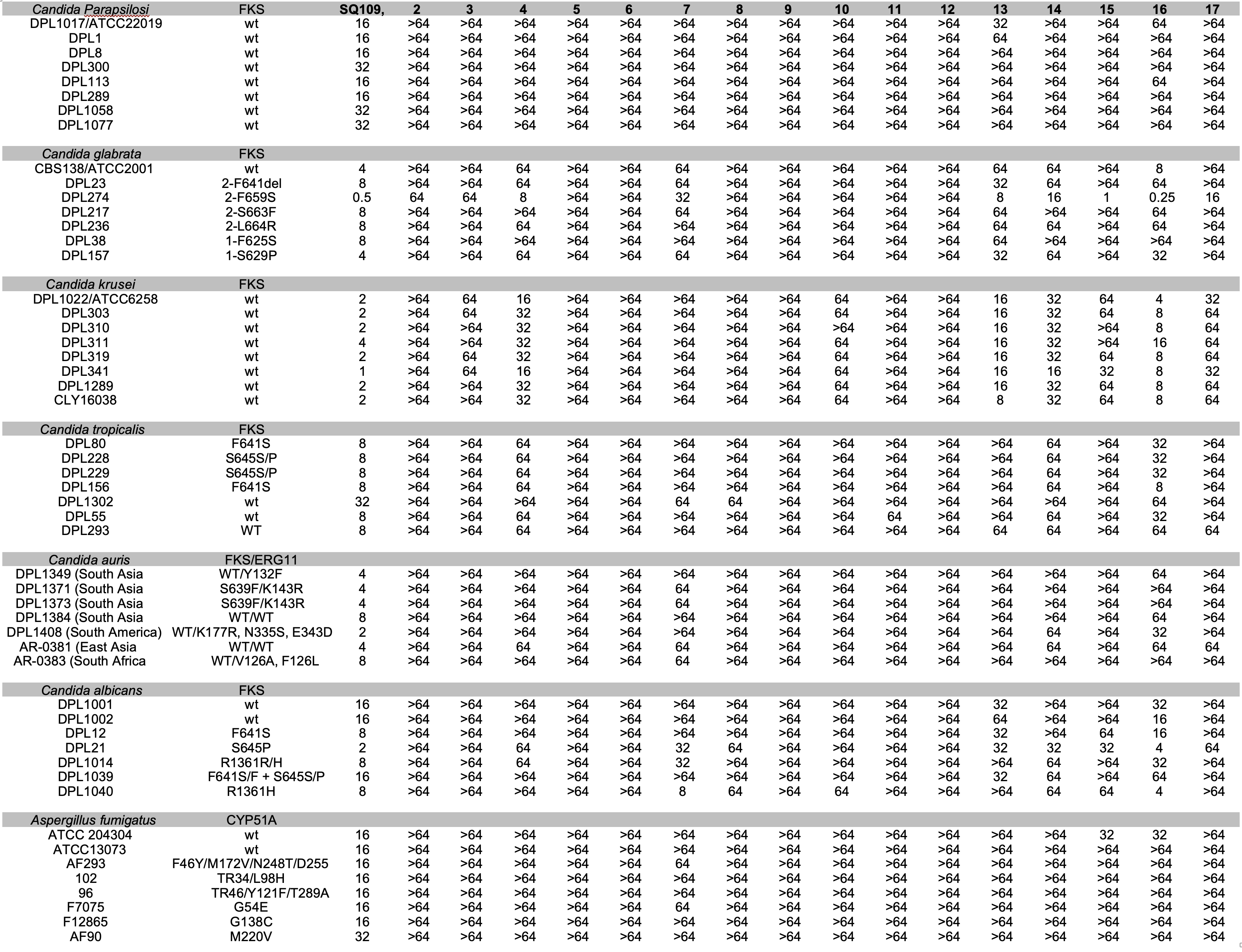

**Table S6. Activity of SQ109 and known/potential metabolites against pathogenic fungi^a^.**

^a^MIC is in mg/mL after 48 h. Abbreviations used are: FKS=1,3-beta-D-glucan-UDP glucosyltransferase; ERG11=lanosterol 14-alpha-demethylase; CYP51A=cytochrome P450 14-alpha sterol demethylase. The mutations in the respective proteins that lead to resistance are indicated.

**Table S7.** ***S. cerevisiae* growth inhibition by SQ109 and SQ109 analogs, and their *E. coli* inverted membrane vesicles (IMV) IC_50_ values.**

| **Compound** | **Structure** | **Sc IC_50_ μM** | **IMV IC_50_ μM** |
| --- | --- | --- | --- |
| **1** | 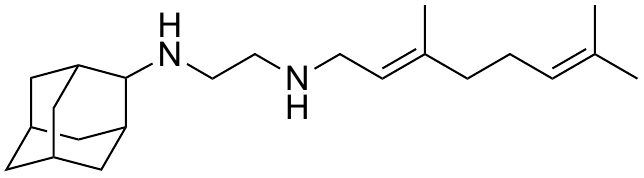 | 1.1 ± 0.02 | 2.3 ± 0.2 |
| **3** | 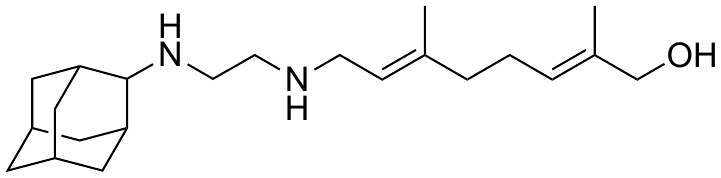 | 166 ± 6 | 17 ± 1 |
| **4** | 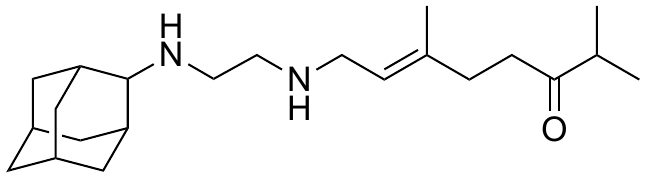 | 136 ± 4 | 15 ± 0.7 |
| **14** | 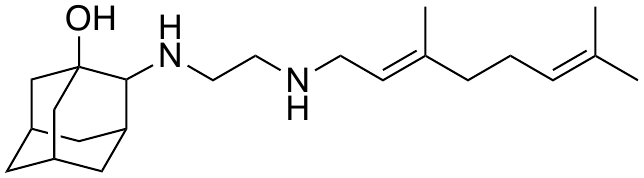 | 51 ± 3 | 9.1 ± 0.03 |
| **15** | 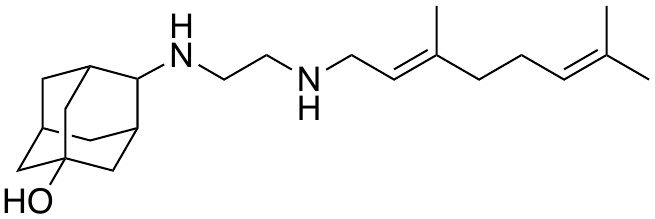 | 14 ± 3 | 18 ± 1 |
| **16** | 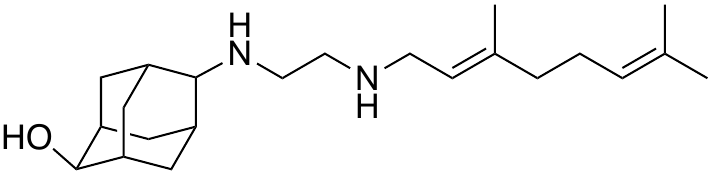 | 13 ± 0.3 | 19 ± 3 |
| **17** | 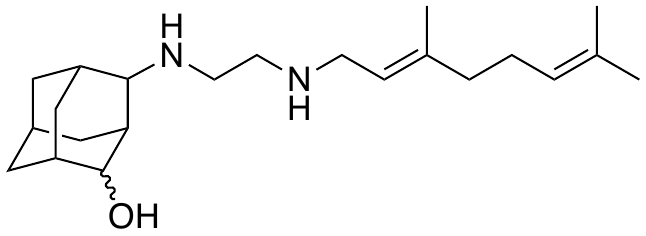 | 59 ± 2 | 12 ± 0.3 |
| **18** | 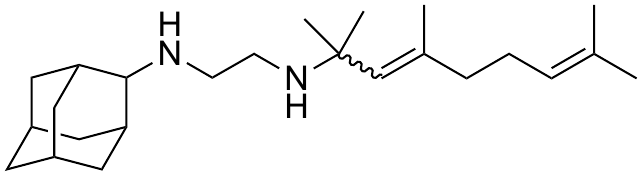 | 8.3 ± 1 | 1.1 ± 0.05 |
| **19** | 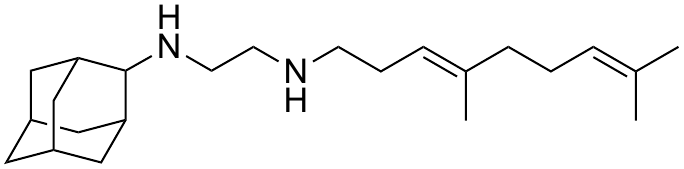 | 3.0 ± 0.4 | 1.9 ± 0.09 |
| **20** | 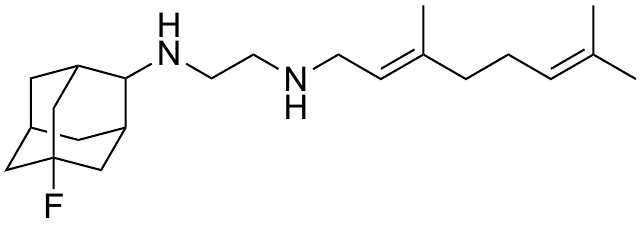 | 8.0 ± 0.4 | 5.5 ± 0.1 |
| **21** | 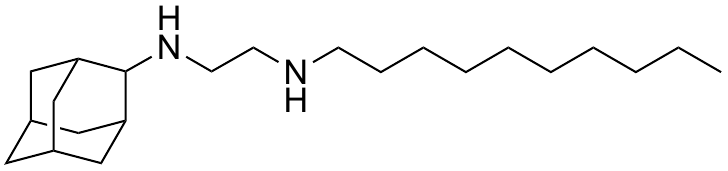 | 7.3 ± 0.05 | 0.87 ± 0.09 |
| **22** | 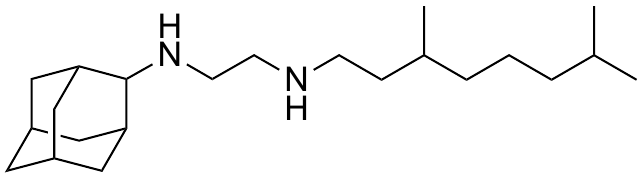 | 7.1 ± 0.2 | 5.2 ± 0.6 |
| **23** | 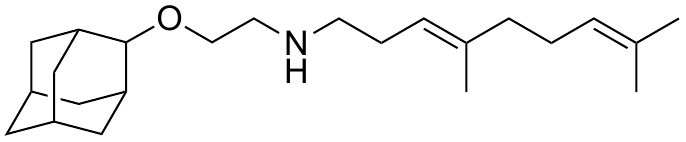 | 3.9 ± 0.3 | 1.2 ± 0.2 |

| **Compound** | | **Structure** | **SMILES** |
| --- | --- | --- | --- |
| **1** | 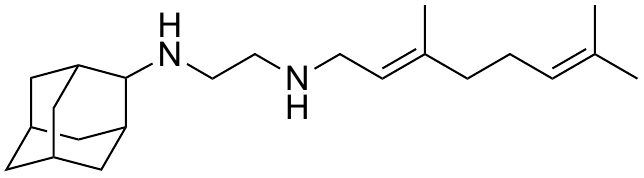 | | C/C=C\CC/C(C)=C/CNCCNC1[C@@H]2C |
| **2** | 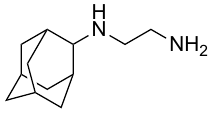 | | NCCNC1[C@@H]2C[C@H](C[C@H]1C3)C[C@H]3C2 |
| **3** | 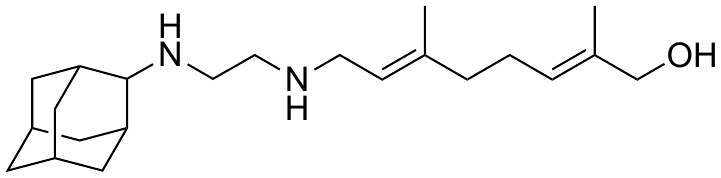 | | C/C(CO)=C\CC/C(C)=C/CNCCNC1[C@@H]2C[C@H](C[C@H]1C3)C[C@H]3C2 |
| **4** | 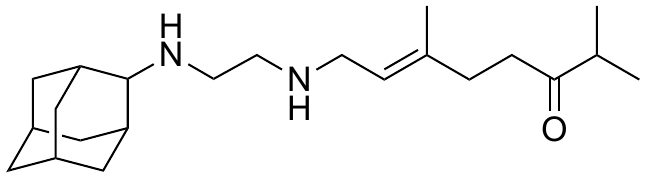 | | CC(C)C(CC/C(C)=C/CNCCNC1[C@@H]2C[C@H](C[C@H]1C3)C[C@H]3C2)=O |
| **5** | 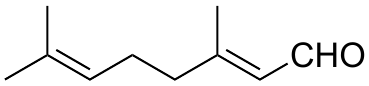 | | C/C(C)=C/CC/C(C)=C/C=O |
| **6** | 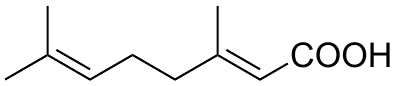 | | C/C(C)=C/CC/C(C)=C/C(O)=O |
| **7** | 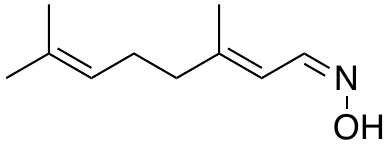 | | C/C(CC/C=C(C)/C)=C\C=N/O |
| **8** | 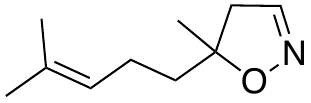 | | C/C(C)=C\CCC1(CC=NO1)C |
| **9** | 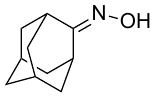 | | O/N=C1[C@@H]2C[C@H](C[C@H]\1C3)C[C@H]3C2 |
| **10** | 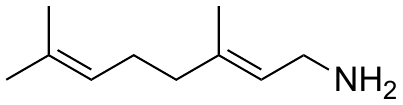 | | NC/C=C/CC/C=C(C)/C |
| **11** | 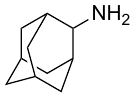 | | NC1[C@@H]2C[C@H](C[C@H]1C3)C[C@H]3C2 |
| **12** | 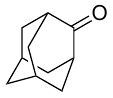 | | O=C1[C@@H]2C[C@H](C[C@H]1C3)C[C@H]3C2 |
| **13** | 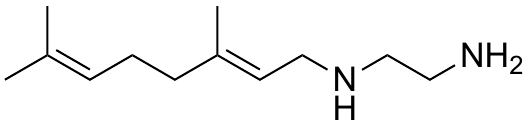 | | NCCNC/C=C(C)/CC/C=C(C)/C |
| **14** | 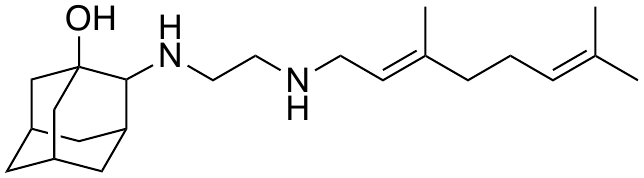 | | C/C(C)=C/CC/C(C)=C/CNCCNC1[C@@H]2C[C@H](C[C@@]1(O)C3)C[C@H]3C2 |
| **15** | 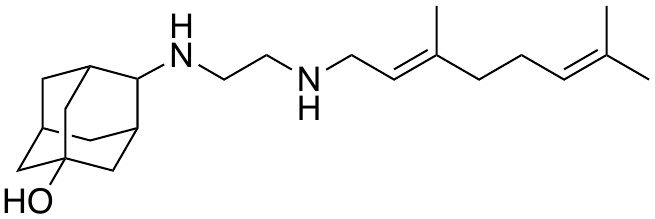 | | C/C(C)=C/CC/C(C)=C/CNCCNC1[C@@H]2C[C@H](C[C@H]1C3)C[C@@]3(O)C2 |
| **16** | 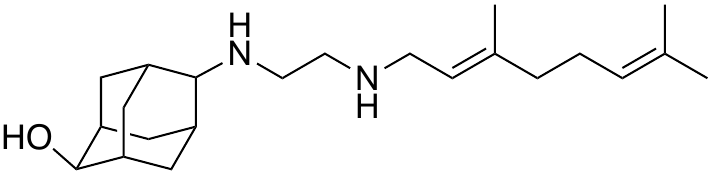 | | C/C(C)=C/CC/C(C)=C/CNCCNC1[C@@H]2C[C@H](C[C@H]1C3)[C@H](O)[C@H]3C2 |
| **17** | 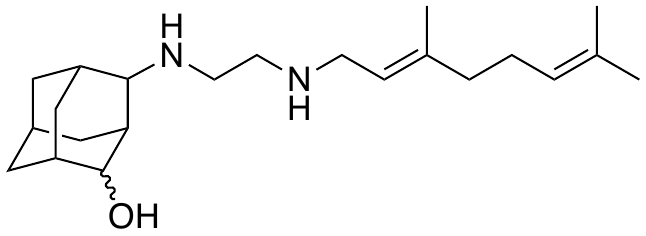 | | C/C(C)=C/CC/C(C)=C/CNCCNC1[C@@H]2C[C@H](C[C@H]1C3)C[C@H]3C2O |
| **18** | 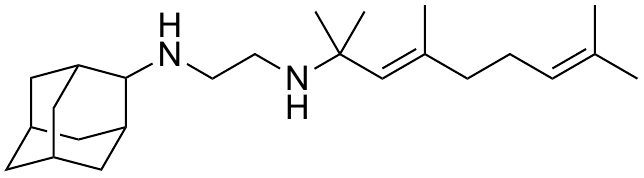 | | C/C(C)=C/CC/C(C)=C/C(C)(C)NCCNC1[C@@H]2C[C@H](C[C@H]1C3)C[C@H]3C2 |
| **19** | 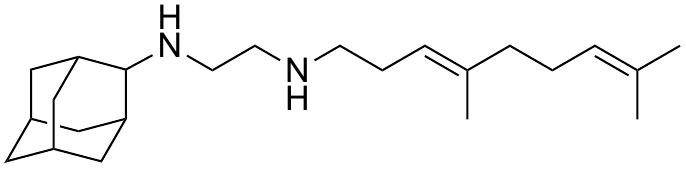 | | C/C(CC/C=C(C)/C)=C\CCNCCNC1[C@@H]2C[C@H](C[C@H]1C3)C[C@H]3C2 |
| **20** | 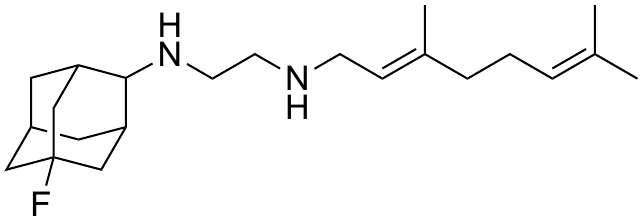 | | C/C(CC/C=C(C)/C)=C\CCNCCNC1[C@@H]2C[C@H](C[C@H]1C3)C[C@@]3(F)C2 |
| **21** | 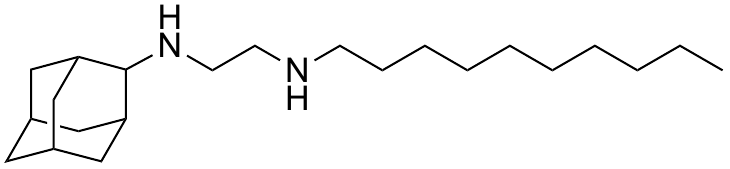 | | CCCCCCCCCCCNCCNC1[C@@H]2C[C@H](C[C@H]1C3)C[C@H]3C2 |
| **22** | 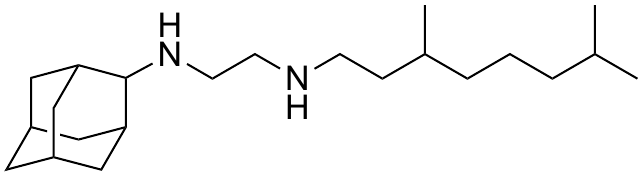 | | CC(C)CCCC(C)CCNCCNC1[C@@H]2C[C@H](C[C@H]1C3)C[C@H]3C2 |
| **23** | 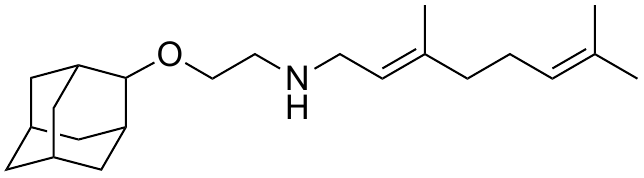 | | C/C(C)=C/CC/C(C)=C/CNCCOC1[C@@H]2C[C@H](C[C@H]1C3)C[C@H]3C2 |
| **24** | 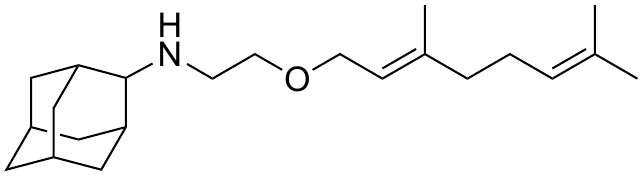 | | C/C(C)=C/CC/C(C)=C/COCCNC1[C@@H]2C[C@H](C[C@H]1C3)C[C@H]3C2 |
